# Parallel signaling through IRE1α and PERK regulates pancreatic neuroendocrine tumor growth and survival

**DOI:** 10.1101/522102

**Authors:** Paul C. Moore, Jenny Y. Qi, Maike Thamsen, Rajarshi Ghosh, Justin Peng, Micah J. Gliedt, Rosa Meza-Acevedo, Rachel E. Warren, Annie Hiniker, Grace E. Kim, Dustin J. Maly, Bradley J. Backes, Feroz R. Papa, Scott A. Oakes

## Abstract

Master regulators of the unfolded protein response (UPR)—IRE1α and PERK— promote adaptation or apoptosis depending on levels of endoplasmic reticulum (ER) stress. While the UPR is activated in many cancers, its effects on tumor growth remain unclear. Derived from endocrine cells, pancreatic neuroendocrine tumors (PanNETs) universally hypersecrete one or more peptide hormones, likely sensitizing these cells to high ER protein-folding stress. For the nearly 1,500 Americans diagnosed with PanNETs annually, surgery is the only potentially curative treatment; however the five-year survival is extremely low for those who develop metastatic disease. To assess whether targeting the UPR is a viable therapeutic strategy, we analyzed human PanNET samples and found evidence of elevated ER stress and UPR activation. We then used genetic and pharmacologic approaches to modulate IRE1α and PERK in cultured cells and xenograft and spontaneous genetic (RIP-Tag2) mouse models of PanNETs. We found that UPR signaling is optimized for adaptation and that inhibiting either IRE1α or PERK leads to hyperactivation and apoptotic signaling through the reciprocal arm, thereby halting tumor growth and survival. Our results provide a strong rationale for therapeutically targeting the UPR in PanNETs and other cancers experiencing elevated ER stress.

**Significance:** The unfolded protein response (UPR) is upregulated in human pancreatic neuroendocrine tumors and its genetic or pharmacological inhibition significantly reduces tumor growth in preclinical models, providing strong rationale for targeting the UPR in neoplasms with elevated ER stress.

## Introduction

When misfolded proteins in the endoplasmic reticulum (ER) accumulate above a critical threshold, the unfolded protein response (UPR) is initiated to restore homeostasis. The UPR is controlled by three ER transmembrane proteins—inositol-requiring enzyme 1α (IRE1α/ERN1), PKR-like ER kinase (PERK/EIF2AK3), and activating transcription factor 6 (ATF6)—that detect misfolded proteins, expand ER protein folding capacity and decrease protein folding demand (1–4).

The most ancient UPR sensor, IRE1α, contains an ER lumenal domain that recognizes unfolded proteins and undergoes dimerization or oligomerization depending on the degree of engagement (5–7). Remediable ER stress causes dimerization and subsequent *trans*-autophosphorylation of the cytosolic kinase domain, which allosterically activates the attached endoribonuclease (RNase) and initiates frame-shift splicing of *XBP1* mRNA. Resulting translation of the XBP1s (s=spliced) transcription factor upregulates genes encoding ER protein-folding and quality control (8, 9). Analogously, recognition of misfolded proteins by the lumenal domain of PERK results in dimerization, *trans*-autophosphorylation and activation of its cytosolic kinase domain (10, 11). PERK then phosphorylates eIF2α, which downregulates Cap-dependent translation, and reduces ER protein load (12, 13). Concurrently, transcripts with upstream open reading frames (uORFs), such as Activating Transcription Factor 4 (ATF4), are preferentially translated and further promote ER protein folding and quality control (14). Finally, during ER stress ATF6 translocates to the Golgi and is cleaved to release its N-terminal transcription factor domain, which upregulates ER quality control factors (15). Collectively, remediable UPR signaling induces an “*Adaptive (A)-UPR*,” which restores ER homeostasis (16).

However, under sustained, irremediable ER stress, the UPR regulators switch from pro-homeostatic to pro-apoptotic outputs (17). High-level kinase autophosphorylation causes IRE1α oligomerization and relaxed RNase specificity, leading to *R*egulated *I*RE1-*D*ependent *D*ecay (RIDD) of ER-localized mRNAs that encode secretory proteins and essential components of the protein-folding machinery (18, 19). This activity not only results in active deterioration of ER function (18), but also degrades select microRNA precursors to upregulate key apoptotic signals, including Thioredoxin-Interacting Protein (TXNIP) (1, 20). Similarly, prolonged PERK activation results in upregulation of the transcription factor CHOP/GADD153, which attenuates expression of anti-apoptotic BCL-2 and increases expression of pro-apoptotic BCL2 family proteins to promote cell death (21, 22). Finally, ATF6 has been reported to reduce expression of the anti-apoptotic protein Mcl-1 and upregulate CHOP (23, 24). Unmitigated ER stress thus culminates in a “*Terminal (T)-UPR*,” whereby adaptive signaling through XBP1s, ATF4 and ATF6 are eclipsed by pro-apoptotic signals (Fig 1A).

**Figure 1.**
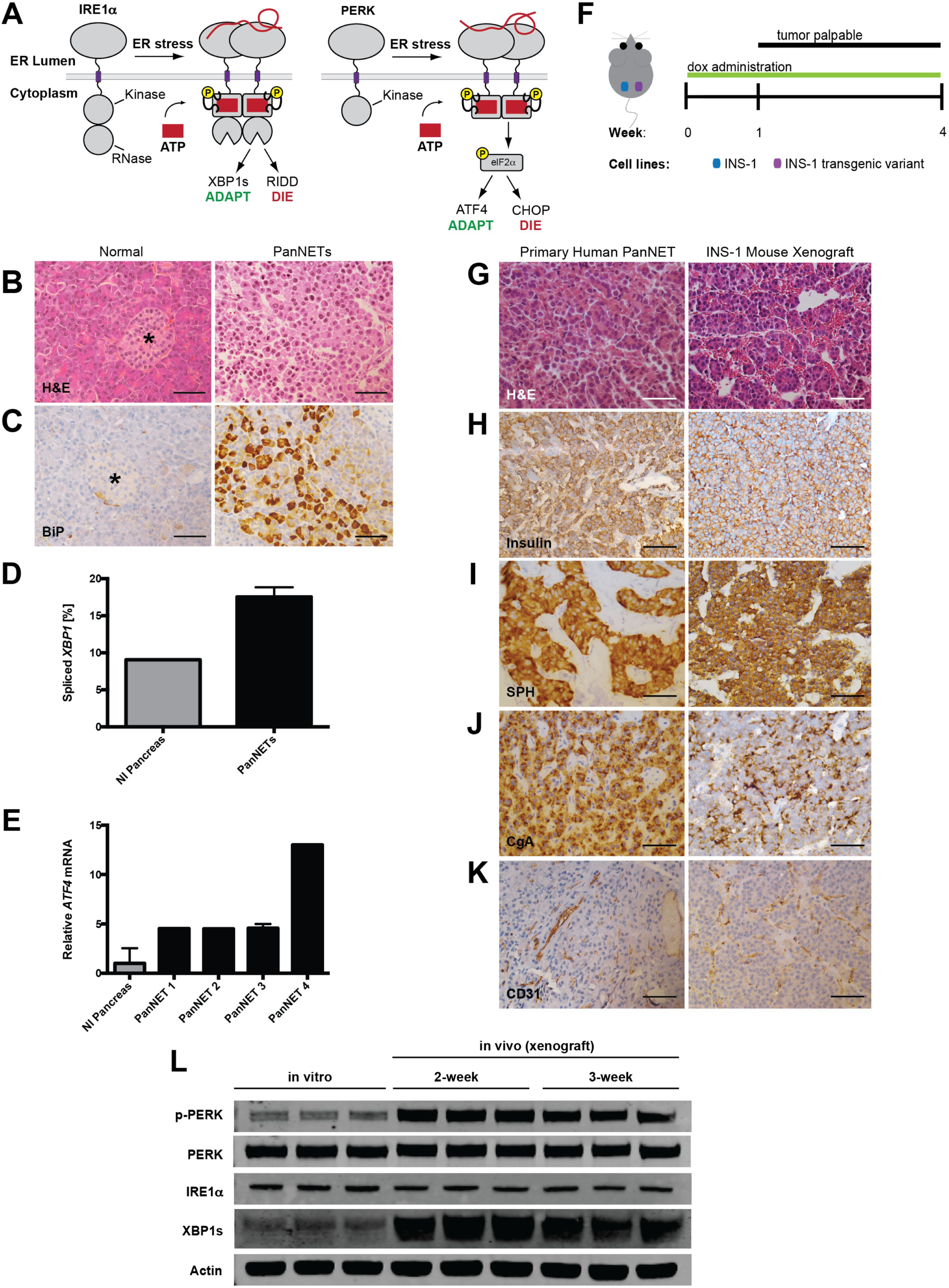
PanNETs show evidence of ER stress and UPR activation. **A**, In response to the accumulation of misfolded proteins in the ER, IRE1α and PERK homodimerize and signal an adaptive stress response through splicing of *Xbp1* and phosphorylation of eIF2α, respectively. However, under sustained ER stress, these pathways promote apoptosis through RIDD and upregulation of pro-apoptotic CHOP. **B-C**, Representative (**B**) H&E and (**C**) BiP/GRP78 IHC on normal pancreas and primary human PanNET. Star indicates islet of Langerhans (scale bars, 50 μm). **D-E**, (**D**) Percent *XBP1* splicing and (**E**) relative *ATF4* mRNA expression from normal human pancreas and four primary human PanNETs. Technical replicate error bars shown in E. **F**, PanNET xenograft experimental setup. INS-1 cells (control vs. transgenic variant) injected s.c. in bilateral flanks of NSG mice. Tumors become palpable by ∼10 d; mice are sacrificed at 4 weeks post-injection. **G-K**, IHC of human PanNETs and INS-1 mouse xenografts stained with the indicated antibodies (CgA=chromogranin A, SPH=synaptophysin; scale bars, 50 μm). **L**, INS-1 cells were grown in tissue culture (*in vitro*) or as xenografts (*in vivo*) in NSG mice for 2 or 3 weeks. Three unique replicates of each condition were harvested and analyzed by immunoblotting with the indicated antibodies. Quantified in Fig. S1B and C.

Elevated ER stress and UPR activity have been documented in various solid cancers, such as glioblastoma and carcinomas of the breast, stomach, esophagus, and liver (25–29). However, whether UPR signaling in these various neoplasms ultimately inhibits or promotes tumor growth remains an area of intense debate (30–34). We speculated that pancreatic neuroendocrine tumors (PanNETs) might be particularly sensitive to protein-folding stress. Derived from endocrine cells, PanNETs universally *hyper*-secrete one or more polypeptide hormones such as insulin or glucagon (35, 36). Moreover, the development and maintenance of pancreatic neuroendocrine cells is greatly impacted by genetic loss of the UPR in mice and humans (37–39).

Here we show that the Adaptive-UPR, especially through the IRE1α and PERK arms, is strongly upregulated in human PanNET samples and in two distinct murine models. Using genetic tools, we discovered that disruption of the Adaptive-UPR or activation of the Terminal-UPR is detrimental to PanNET growth and survival *in vivo*. Likewise, administration of highly selective IRE1α and PERK kinase inhibitors in two murine preclinical PanNET models phenocopies the antitumor effects of genetic deletion. Specifically, inhibiting IRE1α or PERK exacerbates ER stress and leads to apoptotic signaling through the reciprocal UPR branch. Critically, pharmacological targeting of IRE1α increased life expectancy without deleterious effects on animal health, highlighting its therapeutic potential in PanNETs and other ER stress-sensitive cancers.

## Materials and Methods

### Human samples

We obtained 6 formalin-fixed paraffin-embedded (FFPE) samples of de-identified primary human PanNETs, 4 frozen human PanNET samples for RNA analysis, and matched normal pancreata from the UCSF Department of Pathology, IRB protocol number 13-11606

### Tissue culture

Generation of isogenic, stable INS-1 lines was performed as previously described (18). Mycoplasma testing was performed on parent stocks with the MycoAlert Mycoplasma Detection Kit (Lonza) and cells were verified to be mycoplasma free. Cells were cultured in RPMI-1640 media supplemented with 10% tetracycline-free FBS (Gemini Bioproducts 100-800), 10 mM HEPES, 2 mM Glutamine, 110 μg/mL Na Pyr, 100 U/mL penicillin, 100 ug/mL streptomycin (UCSF Cell Culture Facility) and 100 μM βME (Bio Rad 161-0710). All cell stocks were used for ≤ 16 passages. Brefeldin A (9972S) was purchased from Cell Signaling Technology; tunicamycin (T7765), thapsigargin (T9033) and doxycycline (D9891) were purchased from Sigma-Aldrich.

### Small molecules

KIRA8 and was synthesized in-house and purified by reverse phase chromatography (HPLC). The purity of KIRA8 was determined with two analytical RP-HPLC methods, using a Varian Microsorb-MV 100-5 C18 column (4.6 mm x 150 mm), and eluted with either H2O/CH3CN or H2O/ MeOH gradient solvent systems (+0.05% TFA) run over 30 min. Products were detected by UV at 254 nm. KIRA8 was found to be >95% pure in both solvent systems. GSK2656157 (GSK-PKI) was purchased at >98% purity from Advanced Chemblocks Inc. (Burlingame, CA). For use in tissue culture, KIRA8 and GSK-PKI stock solutions were prepared by dissolving in DMSO at a concentration of 20 mM. For use in animal studies, KIRA8 was dissolved in a vehicle solution of 3% ethanol, 7% Tween-80, 1.2% ddH_2_O, and 88.8% of 0.85% W/V saline at a working concentration of 10 mg/ml; GSK-PKI was dissolved in 5% 1-methyl-2-pyrrolidinone (NMP), 5% Kolliphor HS 15 (Solutol), and 90% of 20% W/V (2-Hydroxypropyl)-β-cyclodextrin (HP-β-CD) at a working concentration of 10 mg/mL.

### CRISPR/Cas9

Guide RNAs were designed using the Zhang Lab’s Optimized Design Tool (crispr.mit.edu) and targeted to the 5’ end of each gene to create random insertions/deletions (indels) upstream of key structural and functional domains. For each gRNA, forward and reverse oligonucleotides were purchased from Integrated DNA Technologies (Table S1), annealed, and ligated into the pSpCas9(BB)-2A-EGFP vector (pX458; a gift from Feng Zhang; Addgene Plasmid #48138) at the BbsI cloning site. The resulting plasmids were transfected into INS-1 cells using Lipofectamine 2000 (Thermo Fisher Scientific); a BD FACSAria II (BD Biosciences) was used to subsequently single-cell sort EGFP-positive cells and establish clonal lines. Clones were screened for knockout of target genes by Western Blot and/or by allelic sequencing with custom primers (Integrated DNA Technologies; Table S2) after processing genomic DNA with the KAPA Mouse Genotyping Kit (KAPA Biosystems KK7352) and TOPO TA Cloning Kit (Thermo Fisher Scientific K457501).

### Immunoblot and antibodies

For protein analysis, cells were lysed in T-PER buffer (Thermo Fisher Scientific 78510) plus phosphatase inhibitor cocktail (Cell Signaling Technology #5872S). Protein concentration was determined using Pierce BCA Protein Assay (Thermo Fisher Scientific 23225). Western blots were performed using 10% and 4-12% gradient Bis-Tris precast gels (NuPage) on Invitrogen XCell SureLock Mini-Cell modules. Gels were run using 2-(N-morpholino)ethanesulfonic acid (MES) buffer (Invitrogen NP000202) and transferred onto nitrocellulose transfer membrane using an XCell II Blot Module or iBlot Dry Blotting System (Thermo Fisher Scientific). Antibody binding was visualized on CL-XPosure film using ECL SuperSignal West Extended Duration Substrate (both from Thermo Fisher Scientific) or using the Odyssey CLx Imaging System (LI-COR Biosciences). Antibodies used: ATF4 (Cell Signaling Technology #11815), BiP (CST #3177), CHOP (CST #2895), eIF2a (CST #9722), phospho-eIF2a (CST #3398), IRE1α (CST #3294), PERK (CST #3192), PERK p-T980 (CST #3179), Spliced XBP-1 (BioLegend #619502), actin (Sigma A5441 1:3000).

### RNA isolation, quantitative real-time PCR, and primers

RNA was isolated from whole cells using either Qiashredder and RNeasy kits (Qiagen) or Trizol (Invitrogen). TissueLyser LT (Qiagen) was used for RNA isolation from tumors. For cDNA synthesis, 500-1000 ng total RNA was reverse transcribed using Superscript II Reverse Transcriptase and Oligo d(T)_16_ primer (Invitrogen). For qPCR, we used Power SYBR Green and the StepOnePlus Real-Time PCR System (Applied Biosystems). qPCR primers are listed in Table S3. Gene expression levels were normalized to Actin.

### *Xbp1* mRNA splicing

RNA was isolated from whole cells or tissue and reverse transcribed as detailed above to obtain total cDNA. Sense (5’-AGGAAACTGAAAAACAGAGTAGCAGC-3’) and antisense (5’-TCCTTCTGGGTAGACCTCTGG-3’) primers were used in a standard GoTaq Green PCR reaction (Promega) to amplify a region spanning the 26-nucleotide intron that includes a single PstI restriction site, which is excised by active IRE1α. The resulting PCR fragments were then digested by PstI (New England Biolabs), resolved on 3% agarose gels, stained with ethidium bromide and quantified by densitometry using ImageJ (U. S. National Institutes of Health).

### Cell growth and apoptosis assays

To measure apoptosis by Annexin V staining, cells were plated in 12-well plates overnight. Cells were then treated as described for indicated times. On the day of analysis, cells were trypsinized, washed in PBS, and resuspended in Annexin V binding buffer (10 mM HEPES 7.4, 140 mM NaCl, 2.5 mM CaCl_2_) with Annexin-V FITC (BD Biosciences 556419). Flow cytometry was performed on a Becton Dickinson LSRFortessa or LSRII flow cytometer. To measure cell proliferation, cells were seeded at 5-10% confluence in 96-well plates, treated as indicated, and assayed using the CellTiter-Glo Luminescent Cell Viability Assay (Promega) according to the manufacturer’s protocol. Luminescence was quantified using a Cytation 5 Cell Imaging Multi-Mode Reader (BioTek).

### Animal studies

All animal studies were reviewed and approved by the UCSF Institutional Animal Care and Use Committee. Animals were maintained in a specific pathogen-free animal facility on a 12hr light–dark cycle at an ambient temperature of 21°C. They were given free access to water and food.

### Xenografts

5-8 week old NOD.Cg-*Prkdc^scid^ Il2rg^tm1Wjl^*/SzJ (NSG, Stock #005557, The Jackson Laboratory) mice were injected subcutaneously with 5 x 10^6^ INS-1 cells, and tumor size was followed for up to 4 weeks. Where indicated, animals were provided doxycycline chow (Envigo TD.09761). For small molecule studies, KIRA8, GSK-PKI or the corresponding vehicle solutions were prepared as described above and delivered daily by intraperitoneal (i.p.) injection.

### RIP-Tag2

Tg(RIP1-Tag)2Dh mice (previously described in (40)) were initially obtained from the Bergers Lab at UCSF and maintained as heterozygotes by breeding wild-type C57BL/6 female mice with hemizygous RIP-Tag2 (RT2) male mice. RT2-positive mice were given supplemental diet with adjusted sucrose starting at 12 weeks (Envigo TD.86489). KIRA8 and GSK-PKI treatments, as detailed above, were initiated at 12 weeks and continued as described.

### Glucose tolerance test, blood collection and protocols

After completion of treatment (14 d), mice were starved for 5 h and blood glucose levels were measured to mark time = 0. Mice were then injected with 1 mg/g D-glucose (Thermo Fisher Scientific A24940-01) and their blood glucose monitored over 180 minutes. To monitor blood glucose levels, a drop of blood was collected from the tail onto OneTouch® Ultra® Blue test strips and measured using the OneTouch® Ultra® 2 Meter (LifeScan). For plasma analysis, blood was collected by retro-orbital bleed into EDTA-coated tubes (BD #365974) and analyzed using Amylase Activity Assay Kit (Sigma MAK009-1KT) and Lipase Acitvity Assay Kit (MAK046-1KT) according to the manufacturer’s protocols.

### Histology and immunostaining

Samples were fixed in 4% buffered formaldehyde for 24 h, washed in PBS, transferred into 70% EtOH in ddH_2_O, and then embedded in paraffin and sectioned (5mm thickness) using a Leica RM2255 rotary microtome, by the Brain Tumor Research Center (BTRC) Histology Core, or by the Biobank and Tissue Technology Lab at UCSF. Hematoxylin and eosin staining was performed using standard methods. Stained slides were imaged using an Aperio AT2 slide scanner and data were processed using QuPath software. Total cell counts, Ki67 stains in all tissues, and cleaved caspase-3 stains in RT2 tumors were automated in a blinded fashion; cleaved caspase-3 stains in INS-1 xenografts were quantified manually in a blinded fashion. Antibodies used for immunohistochemistry: BiP [C50B12] (Cell Signaling Technology #3177, 1:200), CD31 (CST #77699 1:100), Chromogranin A [polyclonal] (Cell Marque, 1:4), Cleaved Caspase-3 (CST #9661 1:200), Insulin (DAKO A0564, 1:200), IRE1α (CST #3294, 1:100), Ki67 (Ventana #790-4286, Undiluted), Myc (Sigma M4439, 1:5000), Synaptophysin [LK2H10 clone] (Cell Marque, 1:100).

### Statistical analysis

Graphs were generated using Prism 6 software and represent the average of at least 3 independent experiments. Data are expressed as means ± SD for bar graphs and dot plots when n ≥ 5; for n<5, error bars were excluded. Box plots show min.,*Q_1_*, median, *Q_3_*, and max. Statistical significance was indicated when *P* < 0.05. Two-tailed Welch’s *t* test used for direct comparison between two groups (equal variance not assumed); 1-way ANOVA with Tukey’s multiple comparison test used for ≥ 3 groups; 2-way ANOVA with Tukey’s or Sidak’s multiple comparison test used when two factors (independent variables) were analyzed; logrank (Mantel-Cox) test used for survival analysis.

## Results

### Primary Human PanNETs and a Xenograft PanNET Model Show Evidence of Elevated ER Stress

We obtained a panel of six human PanNETs (Stage I) and performed immunohistochemistry (IHC) against the ER chaperone BiP/GRP78, which is upregulated by the Adaptive-UPR. We observed markedly higher BiP expression in 5 of the 6 human PanNETs compared with normal pancreas (Fig 1B-C, S1A). Moreover, *XBP1* splicing (IRE1α signaling) and *ATF4* mRNA expression (PERK signaling) were upregulated in human PanNETs compared with normal pancreas (Fig 1D-E). We were unable to assess the activation state of ATF6 because all commercially available antibodies tested showed non-specific staining by immunohistochemistry.

To recapitulate UPR signaling *in vivo*, we employed a PanNET xenograft mouse model using rat insulinoma (INS-1) cells (41, 42). We injected INS-1 cells subcutaneously (s.c.) into the flank of immunodeficient NOD-*scid* IL2Rgamma^null^ (NSG) mice (Fig 1F). Tumors became palpable at 1-2 weeks post-injection, grew locally without metastasizing (Stage I), and closely resembled human PanNETs by histology and IHC for known markers (Fig 1G-J). Moreover, CD31 staining demonstrated that the INS-1 xenografts showed similar vascular patterns to human PanNETs (Fig 1K). Notably, compared to INS-1 cells grown *in vitro*, xenograft tumors displayed marked upregulation of IRE1α (XBP1s) and PERK (p-PERK) signaling (Fig 1L; S1B-C).

### Manipulation of IRE1α Adaptive vs. Apoptotic Signaling Determines Growth of INS-1 Xenograft Tumors

We previously engineered transgenic INS-1 cell lines stably integrated with doxycycline (Dox)-inducible constructs of myc-tagged IRE1α (18, 26). *In vitro*, Dox induces supraphysiological production of transgenic IRE1α, which oligomerizes through mass action to induce *Xbp1* splicing, ER-localized mRNA decay, and a Terminal-UPR (18). To test the effects of IRE1α hyperactivation *in vivo*, we injected INS-1::Vector and INS-1::Myc-IRE1α cells into the flanks of NSG mice and provided either regular or Dox chow (Fig 1F). To study long- and short-term effects of IRE1α expression, Dox-induction was provided either for the entire 4-week duration of tumor growth or for a 48-hr interval two weeks post tumor cell injection. Dox chow alone had no effect on the size of control INS-1 tumors over a 4-week time course; in contrast, Dox-induced overexpression of IRE1α markedly reduced tumor mass to <30% of control tumors (Fig 2A-C, S2A) and led to significant increases in both adaptive (*Xbp1s*; Fig 2D, S2B) and apoptotic/RIDD outputs (elevated *Txnip*; decreased *Ins1* and *Ins2*; Fig 2E-G, S2C-D). Even short-term expression of IRE1α was sufficient to induce robust apoptosis as visualized by cleaved caspase-3 staining (Fig 2H-I, S2E and F). Interestingly, IRE1α overexpression resulted in some degree of PERK phosphorylation, perhaps through crowding effects at the ER membrane. However, this was not associated with activation of the downstream effector ATF4 or detectable levels of pro-apoptotic CHOP (Fig S2G-H).

**Figure 2.**
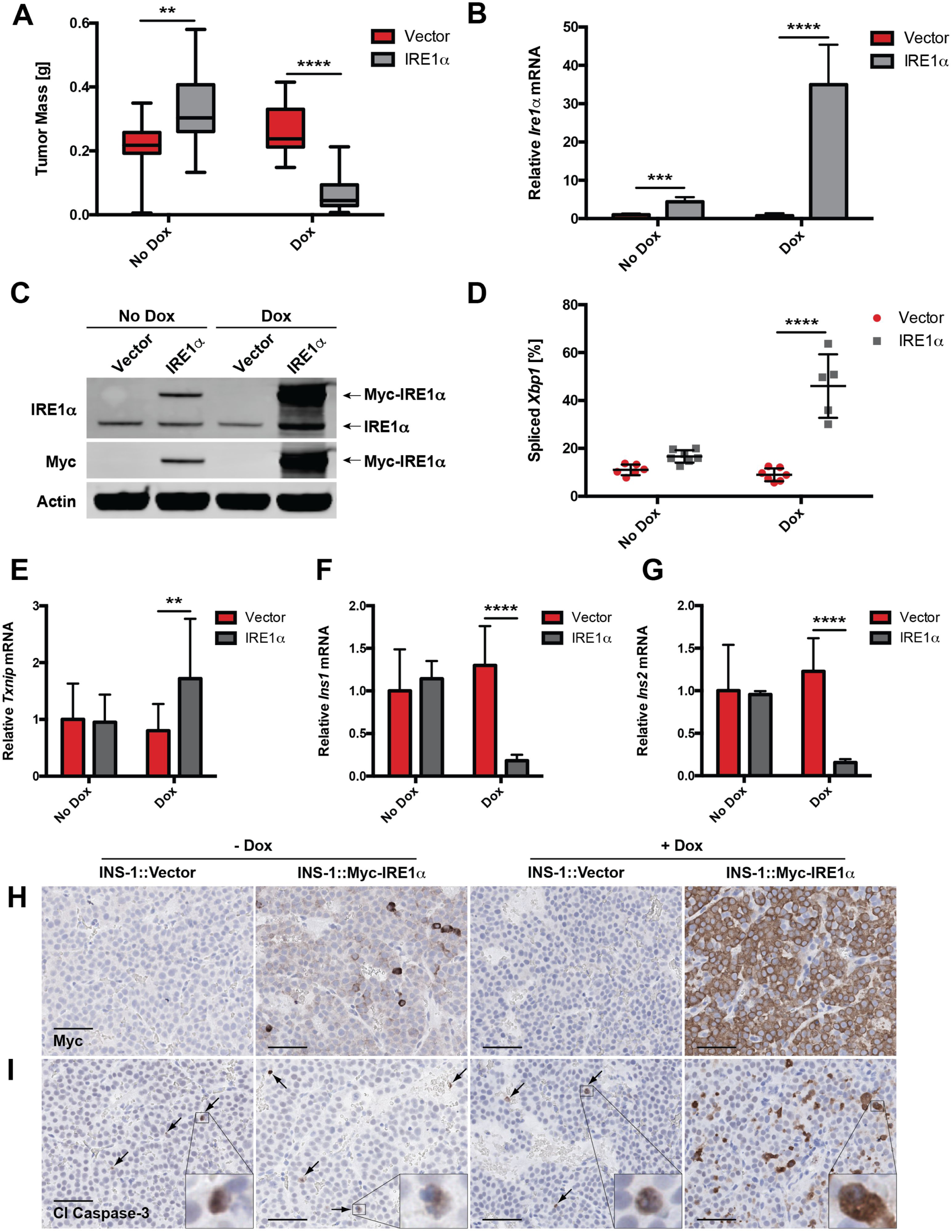
Manipulation of IRE1α adaptive vs. apoptotic signaling determines growth of INS-1 xenograft tumors. **A**, NSG mice were injected s.c. with INS-1 (vector) cells or INS-1 cells carrying a Dox-inducible, myc-tagged Ire1α gene (INS-1::Ire1α). Mice were fed regular or Dox chow as shown in Figure 1F, and tumors were harvested and weighed after 4 weeks (n ≥ 13). **B**, NSG mice developed INS-1::Vector or INS-1::Ire1α tumors for 14 d before administration of regular or Dox chow for 96 h. Tumors were harvested, and *Ire1α* mRNA levels were quantified (n ≥ 5). **C**, Expression of transgenic myc-tagged IRE1α in tumors treated +/-Dox for 96 h at 2 weeks post-injection. Immunoblotted with the indicated antibodies; endogenous and myc-tagged IRE1α species indicated with arrows. **D**, Percent spliced *Xbp1* in INS-1 xenograft tumors treated +/-Dox for 96 h at 2 weeks post-injection (n ≥ 5). **E**, mRNA levels of *Txnip* at 4 weeks post-injection (n ≥ 13). **F-G**, mRNA levels of (**F**) *Ins1* and (**G**) *Ins2* in INS-1 xenograft tumors treated +/-Dox for 96 h at 2 weeks post-injection (n ≥ 5). **H-I**, IHC for (**H**) myc and (**I**) cleaved caspase-3 in INS-1 xenograft tumors treated +/-Dox for 96 h at 2 weeks post-injection (scale bars, 50 μm). *\*\*P<*0.01, \*\*\**P*<0.001, \*\*\*\**P*<0.0001 (2-way ANOVA, Tukey test).

Intriguingly, INS-1::IRE1α tumors were consistently larger than the INS-1 vector controls in the absence of Dox (Fig 2A). The most likely explanation is that leaky expression of the *Ire1α* transgene (Fig 2B-C, S2A and E) promoted a modest increase in *Xbp1* splicing without triggering IRE1α’s apoptotic outputs, as seen in *vitro* (Fig 2D-I, S2B-D; (18)). These results support the notion that the balance of adaptive and apoptotic signals from IRE1α is optimized by PanNETs to favor growth and avert cell death.

### CRISPR/Cas9-mediated Knockout of IRE1α or PERK Pathway Dramatically Decreases INS-1 Tumor Burden

Just as hyperactivation of IRE1α impairs tumor development, we reasoned that inhibition of UPR pathways would also unbalance UPR signaling and impede tumorigenesis. To initially test this concept, we used CRISPR/Cas9 gene editing to functionally knock out *Ire1α*, *Xbp1*, and *Perk* in INS-1 cells (Fig 3A; S3A). Interestingly, proliferation and survival rates of cultured INS-1 KO lines were not diminished in comparison to the parental INS-1 cells (Fig 3B), suggesting that full UPR functionality is dispensable *in vitro*. We next injected the INS-1 KO cell lines into NSG mice and allowed tumors to develop for 4 weeks. While scrambled (Scr) CRISPR/Cas9 controls achieved a tumor mass no different from that of the parental INS-1 cells (Fig S3B), all UPR KO clones showed markedly impaired tumor growth, in some cases attaining only ∼10% of the mass of control tumors (Fig 3C-H). This correlated with a >65% reduction of actively proliferating cells, as determined by Ki67 staining (Fig 3I-J). Neither differences in apoptosis nor evidence of significant signaling compensation in IRE1α and PERK KOs were detected at the 4-week endpoint of the experiment (Fig S3C-E).

**Figure 3.**
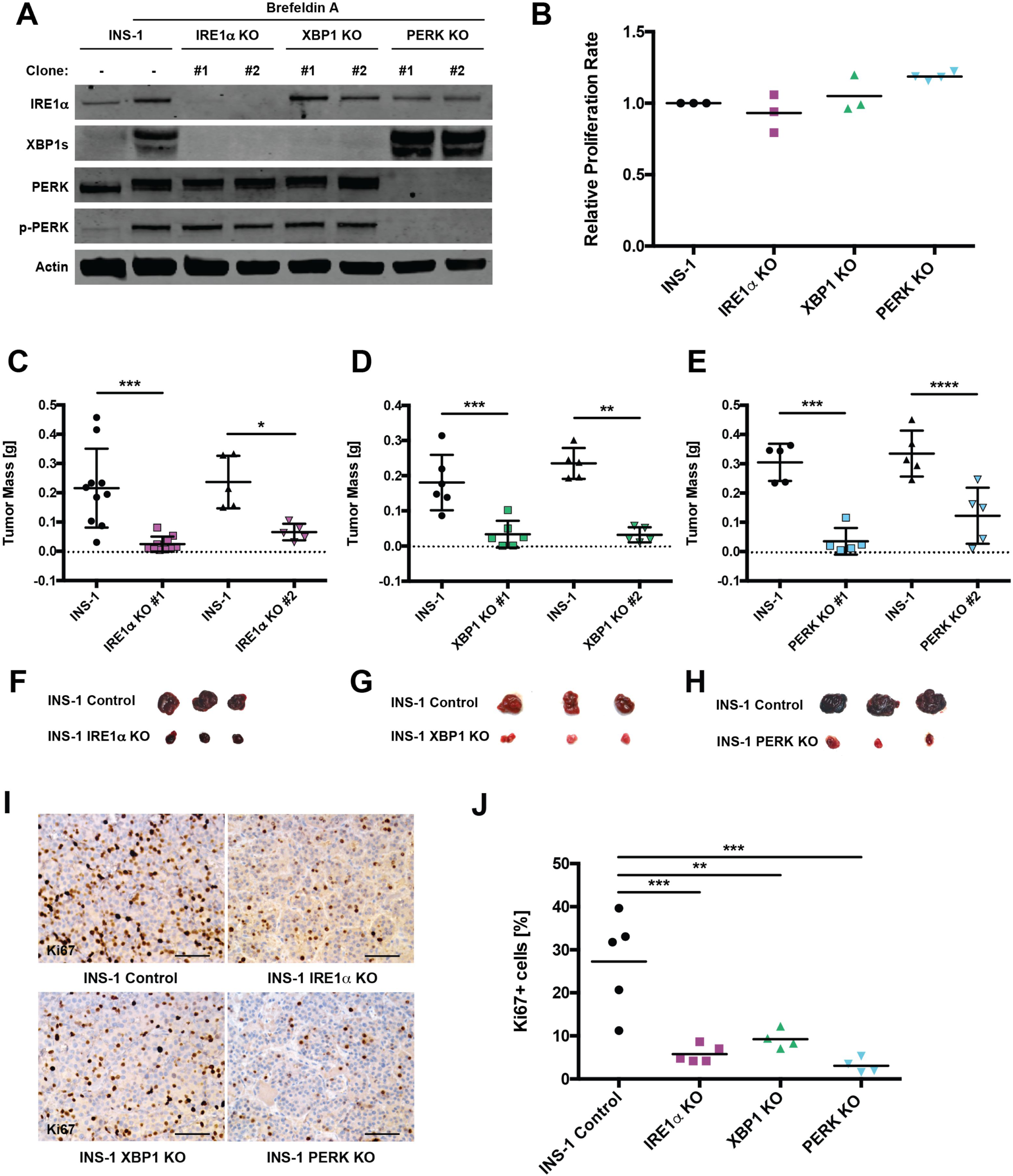
CRISPR/Cas9-mediated knockout of IRE1α or PERK pathways dramatically decreases INS-1 tumor burden. **A**, Cultured INS-1 (control) cells and the indicated CRISPR/Cas9 KO clones were treated +/-0.625 μg/mL Brefeldin A (BFA) for 3 h prior to harvest to induce ER stress and then immunoblotted with the indicated antibodies. **B**, INS-1 cells and the indicated KO lines were subjected to the CellTiter-Glo luminescence-based proliferation assay at 2 and 6 d after seeding. Fold change in luminescence over 96 h was calculated for each KO line and normalized to INS-1 control (n ≥ 3). **C-E**, NSG mice were injected s.c. with INS-1 control and one of two unique (**C**) IRE1α, (**D**) XBP1 or (**E**) PERK KO clones. Resulting tumors were harvested and weighed 4 weeks post-injection (n ≥ 5). **F-H**, Photos of three representative control and (**F**) IRE1α, (**G**) XBP1 or (**H**) PER tumors from C-E. **I**, Representative IHC for Ki67 from control and indicated KO tumors 4 weeks post-injection (scale bars, 50 μm). **J**, Quantification of Ki67 staining in I (n ≥ 4). \**P*<0.05, \*\**P*<0.01, \*\*\**P*<0.001, \*\*\*\**P*<0.0001, \*\*\*\*\**P*<0.00001 (1-way ANOVA, Tukey test).

### Pharmacological Inhibitors of IRE1α and PERK Increase Sensitivity to ER Stress-Induced Apoptosis *In Vitro*

Our team developed ATP-competitive IRE1α *K*inase *I*nhibiting *R*Nase *A*ttenuators— *KIRAs*—that bind IRE1α’s kinase domain and allosterically inhibit its RNase (43, 44). We re-synthesized a monoselective IRE1α inhibitor from a recent KIRA series (compound 18) (45), which has recently been renamed KIRA8 (Fig S4A) (46). Administration of KIRA8 to cultured INS-1 cells revealed an IC_50_ of ∼125 nM; doses between 500 nM and 1 μM resulted in near-complete inhibition of *Xbp1* splicing, reversal of RIDD (*Ins1* mRNA decay), and prevention of apoptosis due to supraphysiological IRE1α signaling (Fig S4B-D).

Correspondingly, we obtained a highly selective PERK inhibitor—GSK2656157— referred to here as *GSK*-*P*ERK *K*inase *I*nhibitor (GSK-PKI; Fig S4E) (47). GSK-PKI displayed minimal off-target inhibition against a panel of 300 kinases *in vitro*, including other eIF2α kinases (47). Under conditions of elevated ER stress in cultured INS-1 cells, GSK-PKI also exhibited an IC_50_ of ∼125 nM with near-optimal inhibition of PERK autophosphorylation and CHOP expression between 500 nM and 1 μM (Fig S4F).

Even at doses that essentially eliminate *Xbp1* splicing, extensive exposure of cultured INS-1 cells to KIRA8 neither affected their viability (Fig 4A) nor their proliferation (Fig 4C). This is consistent with the original report (45) and our findings that IRE1α activity is relatively low and dispensable in cultured cells (Fig 1L and 3B). Conversely, GSK-PKI treatment mildly increased apoptosis and dampened growth rate in cultured INS-1 cells (Fig 4B and C). This discrepancy may stem from pathway compensation in PERK KO cells or off-target GSK-PKI effects (48), but the aggregate effect is modest under low ER stress.

**Figure 4.**
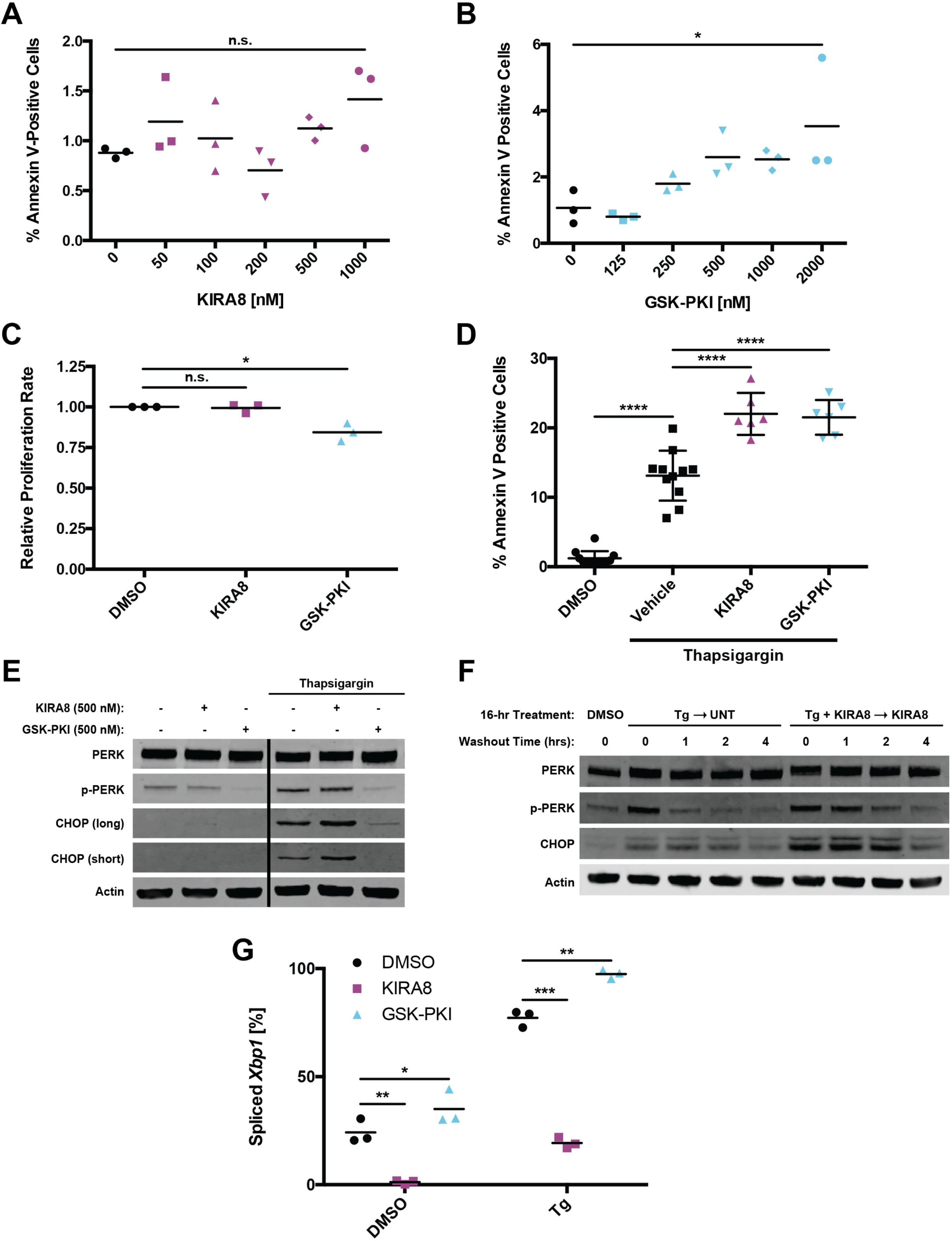
Inhibition of IRE1α or PERK leads to compensatory activation of the other arm and increased sensitivity to ER stress-induced apoptosis. **A-B**, Percentage of cells stained with Annexin V-FITC after five d of treatment with the indicated concentrations of (**A**) KIRA8 or (**B**) GSK-PKI (n = 3). **C**, Normalized fold change in luminescence over 96 h for cells treated with 1 μM KIRA8 or 2 μM GSK-PKI (n = 3). **D**, Percentage of cultured INS-1 cells stained with Annexin V-FITC after 30 h of treatment with the indicated combinations of 31.25 nM thapsigargin, 500 nM KIRA8 and 500 nM GSK-PKI (n ≥ 6). **E**, Cultured INS-1 cells were treated for 20 h with the indicated concentrations of thapsigargin, KIRA8 and GSK-PKI, harvested, and immunoblotted with the indicated antibodies. Both long and short exposures for CHOP are shown. Solid black line indicates excised lane between lanes 3 and 4. **F**, Cultured INS-1 cells were treated concurrently with the indicated combinations of 31.25 nM thapsigargin and 1 μM KIRA8. After 16 h, cells were placed in thapsigargin-free media while KIRA8 treatment was maintained where indicated. Samples were harvested over the indicated time course and immunoblotted with the indicated antibodies. Quantified in Fig. S4H and I. **G**, Percent *Xbp1* splicing from cells treated as in E (n = 3). \**P*<0.05, \*\*\**P*<0.001, n.s. = not significant (1-way ANOVA, Tukey test in A-D; 2-way ANOVA, Tukey test in G).

To mildly induce ER stress and activate both IRE1α and PERK pathways, we used thapsigargin (ER Ca^2+^ pump inhibitor; Fig 4G, S4G) alone or in combination with 500 nM of KIRA8 or GSK-PKI. Under these conditions, the addition of either inhibitor roughly doubled the number of apoptotic cells over 30 hours (h) of treatment (Fig 4D), raising the possibility of compensatory signaling through a parallel pathway. While treatment with KIRA8 did not noticeably affect PERK autophosphorylation, high baseline levels may have masked any subtle changes (Fig 4E). However, KIRA8 increased thapsigargin-induced CHOP expression (Fig 4E) and sustained p-PERK and CHOP levels after thapsigargin washout (Fig 4F; S4H and I). Inversely, treatment with GSK-PKI boosted *Xbp1* splicing, both with and without ER stress induction, indicating compensation through IRE1α (Fig 4G).

### Pharmaceutical Inhibition of IRE1α or PERK Impedes Growth and Induces Apoptosis of INS-1 Tumors

We next tested the effectiveness of pharmacologic UPR inhibitors in the INS-1 xenograft model by administering doses of KIRA8 or GSK-PKI estimated to achieve serum concentrations above IC_50_. Tumors were grown in NSG mice for two weeks, treated for 48 h, harvested, and analyzed for target inhibition by their respective drugs. Dosing with KIRA8 lowered *Xbp1* splicing from a baseline of ∼15% to below 1.5% (Fig 5A; S5A); GSK-PKI treatment reduced PERK autophosphorylation below detectable levels (Fig 5B).

**Figure 5.**
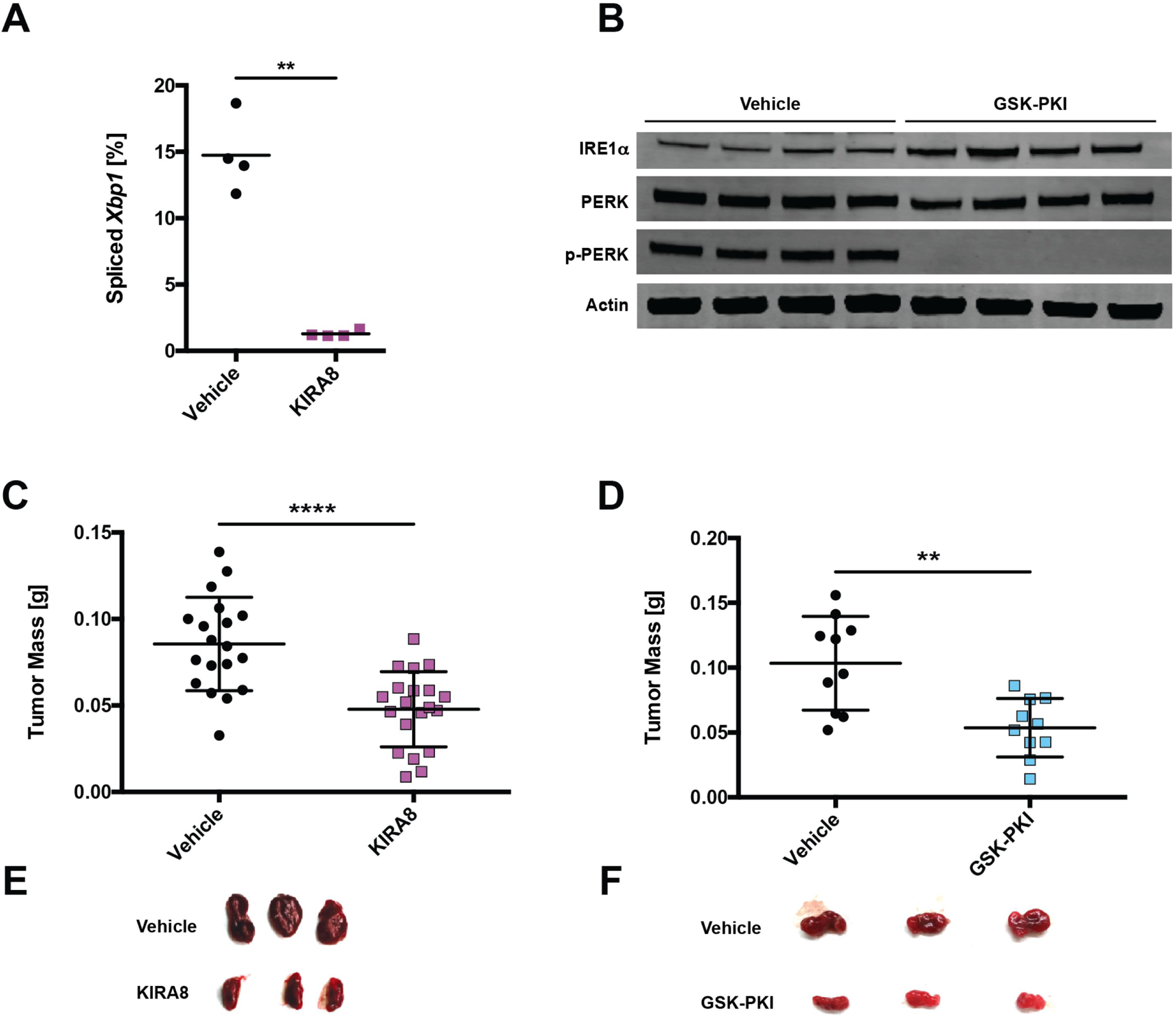
Pharmacological inhibition of Ire1α or Perk decreases INS-1 tumor size. **A**, NSG mice were injected s.c. with INS-1 cells and 2 weeks later injected i.p. daily with vehicle or 50 mg/kg KIRA8. Tumors were harvested after 48 h of vehicle or KIRA8 treatment and subjected to *Xbp1* splicing analysis (n = 4). **B**, NSG mice were injected s.c. with INS-1 cells and 2 weeks later injected i.p. daily with vehicle or 50 mg/kg GSK-PKI. Tumors were harvested after 48 h of vehicle or GSK-PKI treatment and immunoblotted with the indicated antibodies. **C-D**, NSG mice were injected s.c. with INS-1 cells and administered (**C**) 40 mg/kg/d KIRA8 or (**D**) 50 mg/kg/d GSK-PKI. Resulting tumors were harvested and weighed at 3 weeks post-injection (n ≥ 10). **E-F**, Photos of three representative vehicle- and (**E**) KIRA8- or (**F**) GSK-PKI-treated tumors from C and D, respectively. \*\**P*<0.01, \*\*\*\**P*<0.0001 (unpaired, two-tailed *t* tests).

To determine effects on tumor burden, we injected NSG mice with INS-1 cells and the following day began administering daily doses of KIRA8 or GSK-PKI for three weeks prior to harvest. While less dramatic than IRE1α or PERK KO tumors, both KIRA8 and GSK-PKI significantly reduced tumor mass by ∼50% (Fig 5C-F). In contrast to cultured INS-1 cells, INS-1 tumors exhibited markedly decreased Ki67 staining within 48 h of KIRA8 or GSK-PKI administration *in vivo* (Fig 6A-D). More specifically, KIRA8 and GSK-PKI blocked G_1_-S transition by impairing expression of Cyclin E1 and D1, respectively (Fig S5B-C).

**Figure 6.**
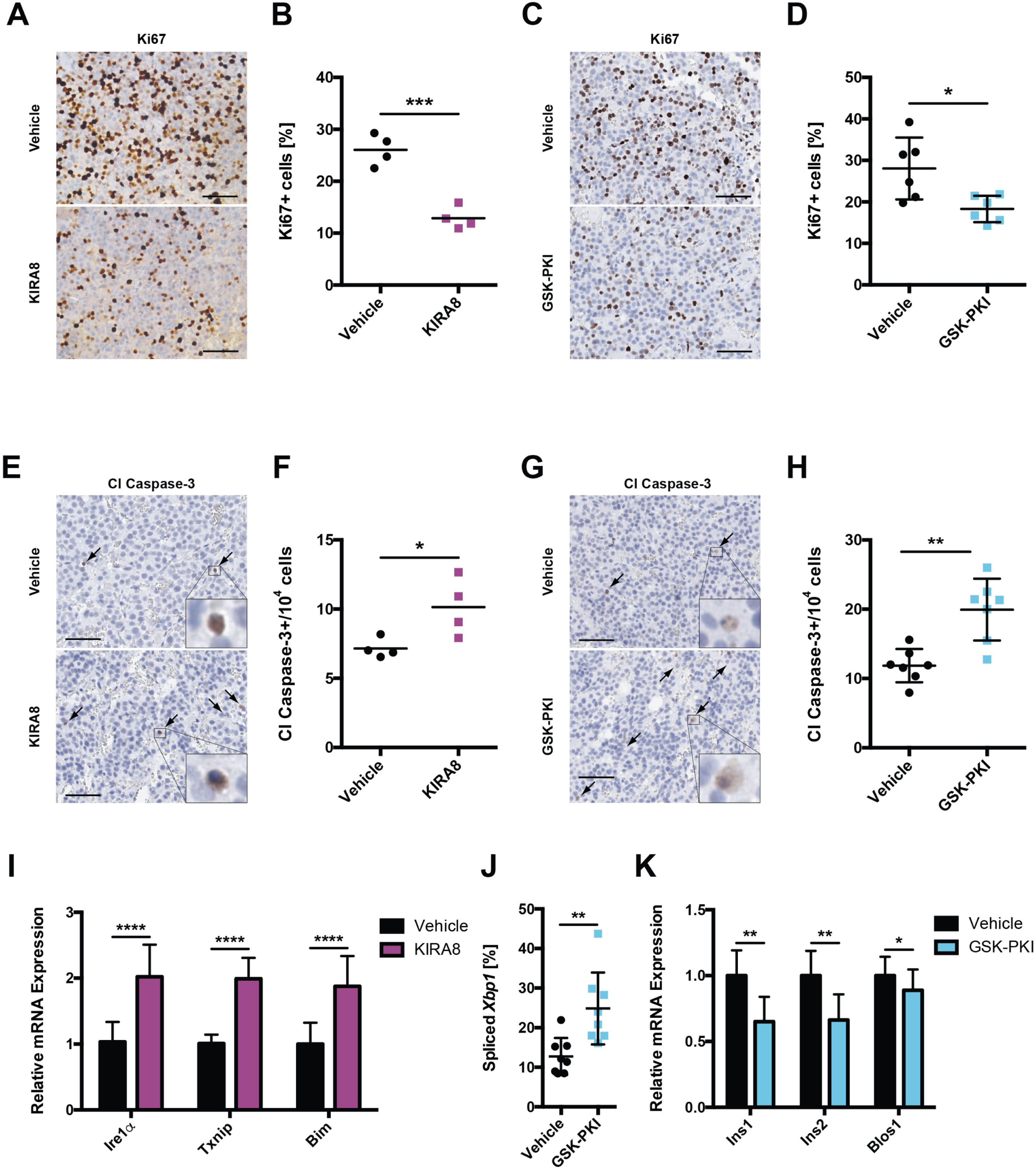
KIRA8 and GSK-PKI induce cell-cycle arrest and apoptosis in INS-1 tumors. **A-B**, Representative IHC for Ki67 in tumors treated for 48 h with (**A**) 50 mg/kg/d KIRA8, (**B**) 50 mg/kg/d GSK-PKI, or the corresponding vehicle (scale bars, 50 μm). **C-D**, Quantification of Ki67 positive cells from A and B, respectively (n ≥ 4). **E-F**, Representative IHC for cleaved caspase-3 in tumors from A and B, respectively (scale bars, 50 μm). **G-H**, Quantification of cleaved caspase-3 positive cells from E and F, respectively (n ≥ 4). **I**, Levels of the indicated mRNAs from tumors in A and E (n ≥ 7). **J**, Percent *Xbp1* splicing in tumors from B and F (n = 8). **K**, mRNA levels of *Ins1, Ins2* and *Blos1* in tumors from B and F (n ≥ 8). \**P*<0.05, \*\**P*<0.01, \*\*\**P*<0.001, \*\*\*\**P*<0.0001, \*\*\*\*\**P*<0.00001 (unpaired, two-tailed *t* tests).

Moreover, *in vivo* administration of either compound increased levels of cleaved caspase-3 roughly 1.5-fold (Fig 6E-H). Due to high basal activation *in vivo*, we were unable to directly detect further increases in p-PERK under KIRA8 treatment (Fig S5D) but did note increased levels of PERK-associated markers for ER stress (*Ire1α*) and apoptosis (*Txnip*, *Bim*; Fig 6I) (1, 49–51). In regard to GSK-PKI, cytotoxicity was associated with increased *Xbp1* splicing (Fig 6J, S5E) and decreases in *Ins1, Ins2, and Blos1* mRNA (Fig 6K), implicating upregulation of adaptive and pro-apoptotic (RIDD) IRE1α signaling, respectively.

### KIRA8 Treatment Decreases Tumor Size and Prolongs Survival in a RIP-Tag2 PanNET Model

We next tested the effects of KIRA8 and GSK-PKI in a second preclinical PanNET model. The RIP-Tag2 (RT2) mouse is a transgenic strain in which viral SV40 large T-antigen (Tag) expression is driven by the β-cell specific rat insulin promoter-1 (RIP) (40), leading to predictable development of islet hyperplasia (5-10 wks), adenomas (10-12 wks), and eventually invasive (12-15 wks) (52)(Fig 7A). Similar to INS-1 xenografts, RT2 tumors also display elevated ER stress compared to wild-type (WT) pancreata. By 10-14 weeks *Xbp1* splicing, IRE1α expression, and PERK phosphorylation are all increased (Fig 7B-C; S6A-B); eIF2α phosphorylation, which integrates multiple stress pathways, is unchanged at the tested time points.

**Figure 7.**
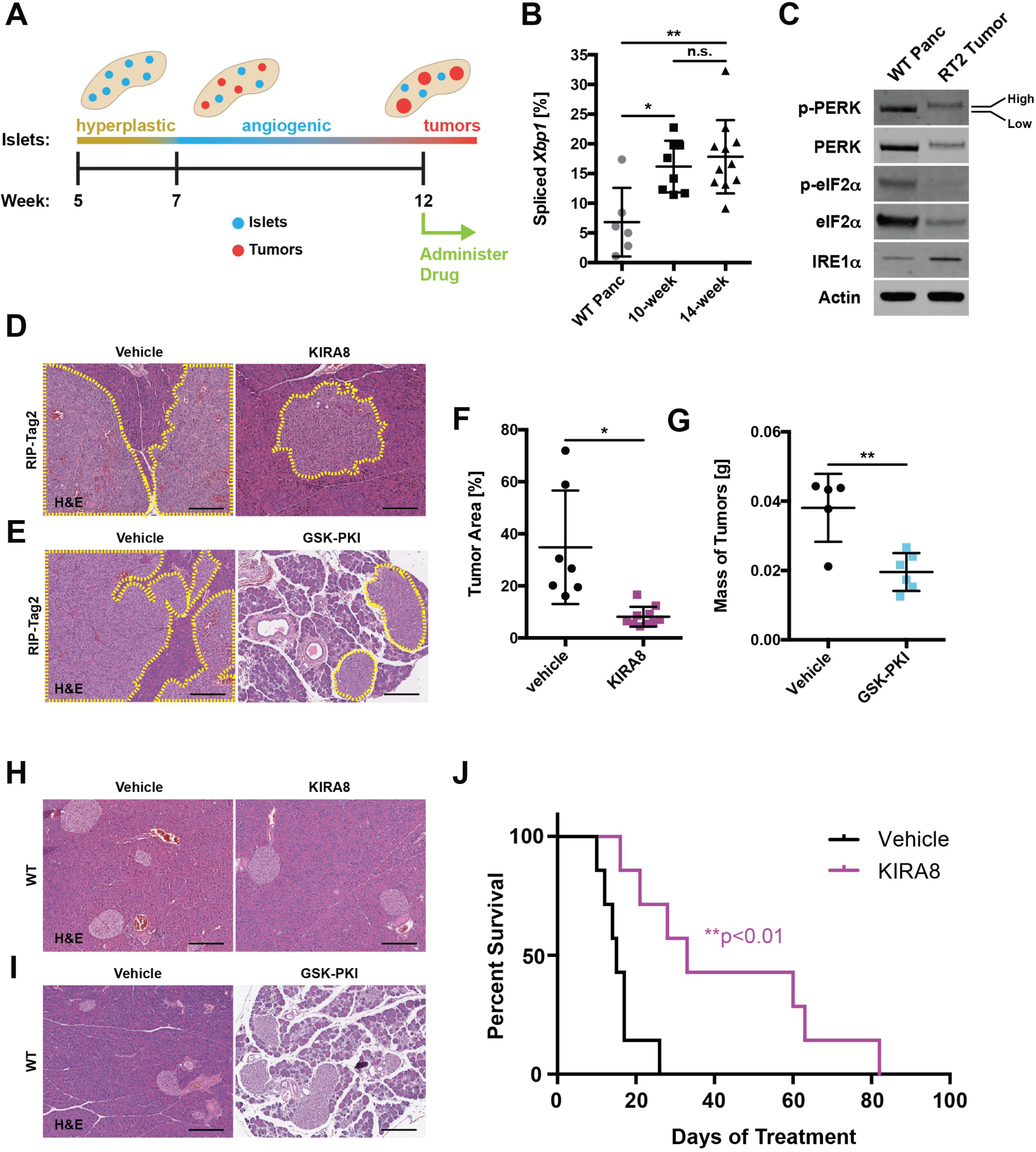
KIRA8 treatment decreases tumor size and prolongs survival in a RIP-Tag2 PanNET model. **A**, Timeline of islet tumorigenesis in the RT2 model. **B**, Percent *Xbp1* splicing in pancreatic tissue taken from 14-week-old WT and PanNET tumors from 10- or 14-week-old RT2 C57BL/6 mice (n ≥ 6). **C**, Tissue from WT and RT2 mice as in B, harvested and immunoblotted with the indicated antibodies. Low and High phosphorylation states of PERK species are indicated. Expanded in Fig. S6A; quantified in S6B. **D-E**, Representative H&E stains of RT2 pancreata from mice treated with (**D**) 50 mg/kg/d KIRA8, (**E**) 50 mg/kg/d GSK-PKI, or the corresponding vehicle beginning at 12 weeks of age and harvested two weeks later (14 d of treatment). Neuroendocrine tissue outlined in yellow (scale bars, 200 μm). **F**, Percent neuroendocrine tissue area in H&E-stained pancreata from RT2 mice treated with vehicle or KIRA8 as in D (n ≥ 7). **G**, Mass of pancreatic tumors harvested from RT2 mice treated with vehicle or GSK-PKI as in D (n ≥ 5). **H-I**, Representative H&E stains of pancreata from 14-week-old WT C57BL/6 mice after 14 d of treatment with (**H**) 50 mg/kg/d KIRA8, (**I**) 50 mg/kg/d GSK-PKI, or the corresponding vehicle (scale bars, 200 μm). **J**, Survival curves of RT2 mice treated with vehicle or 50 mg/kg/d KIRA8 from 12 weeks of age until death (n = 7). \**P*<0.05, \*\**P*<0.01 and n.s. = not significant (1-way ANOVA, Tukey test in B; unpaired, two-tailed *t* tests in F-G; logrank test in J).

RT2 animals were subjected to a tumor regression trial: we began vehicle or drug (KIRA8 or GSK-PKI) administration at 12 weeks of age, when tumors have become invasive (Fig 7A), and continued therapy for 14 days (d). After the 2-week treatment period, we sacrificed the animals and measured tumor burden histologically as a percentage of total pancreas area (Fig 7D-E). In KIRA8-treated animals, *Xbp1* splicing was completely ablated (Fig S6C) and tumor burden was decreased >75% compared with vehicle-treated control animals (Fig 7D and F). There was no visible damage to the surrounding pancreas. As in INS-1 xenografts (Fig 6 and S5), KIRA8 treatment resulted in moderate decrease of Ki67 staining and roughly 2-fold increase of cleaved caspase-3 (Fig S6D-G).

GSK-PKI treatment was associated with widespread degeneration of the exocrine pancreas (Fig 7E), confounding our ability to measure tumor burden histologically; however, direct measurement of tumor masses revealed that GSK-PKI also reduces tumor burden (∼50%) in RT2 animals (Fig 7G). Surprisingly, short-term treatment of RT2 tumors did not reveal changes in Ki67 or cleaved caspase-3 staining despite complete inhibition of PERK activation (Fig S6H-L).

The deleterious impact of GSK-PKI on pancreatic health spurred us to scrutinize the effects of both compounds on WT mice. Twelve-week-old C57BL/6 mice were treated for 14 d with vehicle, KIRA8, or GSK-PKI before sacrificing and removing the pancreas. Weekly blood draws and a terminal glucose tolerance test were performed on a parallel cohort of animals. As expected, KIRA8-treated animals had no discernible damage or loss of pancreas mass (Fig 7H; S7A). Moreover, they recovered normally in a glucose tolerance test and did not display elevated levels of serum amylase or lipase (Fig S7B-D). However, they did experience a ∼10% decline in body mass (Fig S7E). In contrast, GSK-PKI treatment resulted in severe disruption of the exocrine pancreas, a corresponding ∼50% loss of pancreas mass, and a small loss of body mass (Fig 7I; S7F-G. Unexpectedly, they recovered well during a glucose tolerance test and had normal levels of serum amylase and lipase (Fig S7H-J).

Based on these data, we performed a survival study following initiation of KIRA8 treatment at 12 weeks of age. While the vehicle-treated animals lived an average of 17 d after initiation of treatment, the KIRA8 animals survived over twice as long, with several animals surviving over 60 d and one animal up to 82 d (Fig 7J). Because RT2 mice typically did not show overt symptoms prior to death and pancreatic autolysis occurs rapidly, necropsies were not performed. However, all indications are that KIRA8-treated animals eventually succumbed due to PanNET growth similar to the vehicle-treated animals.

## Discussion

Currently, PanNETs are only potentially curable through surgical resection. For the 20-30% of patients diagnosed with metastatic disease, treatment is limited to managing symptoms of hormonal hypersecretion and systemic chemotherapy, to which the tumor invariably develops resistance. The 5-year survival for these patients is as low as 4-25% (53). Treatments with targeted therapies, such as the FDA-approved drugs everolimus (an mTOR inhibitor) and sunitinib (a multi-kinase inhibitor) have a ∼6 month increase in progression-free survival in patients with metastatic PanNETs (54, 55). Recently, peptide receptor radiotherapy (PRRT) has been FDA approved for some patients with metastatic somatostatin receptor (SSTR)-expressing PanNETs, but not all patients respond and the long-term outcomes remain unknown (56).

Notably, professional secretory cells of the endocrine pancreas that PanNETs originate from are critically dependent on the UPR for their development and survival. Generally, deletions of *Perk*, *Ire1α* and *Xbp1* in mice lead to endocrine cell apoptosis and pancreatic dysfunction (37–39). However, it is likely that PanNETs are even more dependent on the UPR as they constitutively hypersecrete one or more hormones (35, 36); some “insulinoma” PanNETs secrete over 10-fold higher amounts of insulin than normal pancreatic β-cells (57). Based on these observations, we predicted that PanNETs would be particularly dependent on the UPR to manage ER stress.

Experiments using inducible IRE1α expression provided the first evidence that the level of UPR activation has important consequences for tumor growth *in vivo*: hyperactivation of IRE1α resulted in massive apoptosis, while moderate expression enhanced adaptive signaling and promoted tumor growth. As such, we hypothesized that downregulation of adaptive UPR signaling would also impede tumor development. Interestingly, neither genetic deletion nor pharmacological blockade of IRE1α or PERK pathways had profound effects on INS-1 growth *in vitro*, consistent with the lack of KIRA8 cytotoxicity previously observed in cultured cancer cell lines (45). While cancer cells face unique challenges to ER proteostasis *in vivo* (e.g., hypoxia, nutrient deprivation), these factors are not mimicked in normal cell culture conditions. Demonstrating this point, we found that INS-1 cells grown in culture have low levels of ER stress and UPR activation compared with the same cells grown in mice as xenografts. By artificially inducing ER stress in cultured INS-1 cells, we increased their dependency on the Adaptive-UPR and sensitized them to KIRA8- and GSK-PKI-induced apoptosis. *In vivo*, IRE1α or PERK inhibition markedly impaired tumor development by decreasing proliferative capacity and inducing apoptosis.

Together, these data provide mechanistic insights into the anti-tumor effects of KIRA8 and GSK-PKI. *In vitro*, inhibiting IRE1α leads to higher sustained activation of PERK and its pro-apoptotic target CHOP. Inversely, PERK inhibition results in IRE1α hyperactivation. In both cases, this leads to loss of an adaptive arm of the UPR, exacerbates ER stress, and shifts the burden to the remaining branch, triggering its apoptotic (Terminal) program. Likewise, *in vivo* inhibiting one arm of the UPR, led to compensatory hyperactivation of the other: IRE1α inhibition by KIRA8 led to transcriptional upregulation of PERK-associated ER stress and apoptotic markers; PERK blockade by GSK-PKI increased IRE1α’s adaptive and apoptotic outputs (RIDD). Although GSK-PKI also inhibits the kinase RIPK1, this is unlikely to contribute to the phenomena observed here as RIPK1 is pro-apoptotic and the drug phenocopies on-target, genetic knockout of *Perk*. Whether ATF6 also experiences compensatory activation is unclear with currently available tools and will require further analysis.

To determine whether our results generalize to other PanNET models, we tested KIRA8 and GSK-PKI in the well-characterized transgenic RT2 model. In use for over 20 years, this model has predicted the clinical efficacy of several compounds that have gone on to FDA approval for PanNETs, including everolimus and sunitinib. In contrast to the INS-1 xenograft model, RT2 tumors arise directly from endogenous pancreatic β-cells and develop in their natural environment. Although both KIRA8 and GSK-PKI are effective at reducing tumor burden in this model, the stark difference in their pancreatic toxicities sets them apart. Within two weeks, daily GSK-PKI administration to WT mice severely damaged and reduced pancreatic mass, similar to results obtained using oral administration of this PERK inhibitor in CD-1 mice (47).

Because genetic deletion of either *Perk* or *Ire1α* can lead to β-cell dysfunction and apoptosis, it was assumed that KIRAs would have similar toxicities to PERK inhibitors. However, inhibiting IRE1α with KIRA8 (or related compound KIRA6) was well tolerated and preserved pancreatic β-cell health in multiple diabetes models (44, 46). Likewise, KIRA8 administration had no noticeable effect on pancreatic mass or histology in WT C57BL/6 or RT2 mice while reducing tumor growth and increasing survival in the RT2 model. This discrepancy in KIRA8 toxicity between tumor and normal pancreatic tissues likely hinges on the post-mitotic nature and low proliferation rate of endogenous pancreatic β-cells. As such, IRE1α emerges as a more attractive therapeutic target than PERK, though more studies are needed to understand the long-term effects of IRE1α inhibition.

Other facets of our data hint at alternative strategies for targeting IRE1α in PanNETs. For one, KIRA8 had a more dramatic effect on tumor burden in the RT2 model than in the INS-1 xenograft model. The difference in immunocompetence between NSG (xenograft) and C57BL/6 mice (RT2 background) may be a decisive factor. For example, recent reports suggest that targeting the IRE1α-XBP1 axis may engage a two-pronged attack that restrains malignant cells while simultaneously eliciting concomitant antitumor immunity (58). The recent explosion in immunotherapy suggests that the immune system has a more complex and widespread role in cancer than initially appreciated, and manipulating the UPR may enhance this approach (59, 60).

Together, our findings reveal many crucial aspects of studying and targeting the UPR in PanNETs. Both the intrinsic secretory nature of PanNETs and their surrounding environments contribute to their level of ER stress, forcing them to lean on the Adaptive-UPR for survival and making them susceptible to pharmacological intervention. Ultimately, optimization of current inhibitors, exploration of combination therapies, and enhancement of UPR activators will diversify our treatment strategies for targeting the UPR in secretory cancers. Moreover, because this approach is not dependent on the presence of a somatic mutation in a UPR component, it may have benefits in many other solid tumors where ER stress is documented.

## Acknowledgements

We thank the Tlsty Lab for NSG mice, IHC protocols, and equipment; Dr. Rushika Perera for assistance with RT2 tumor analysis; the Cyster lab for their microscope; the Goga and Maltepe Labs for providing incubators; the Goga and Debnath Labs for antibodies; the Hebrok Lab for islet histology expertise; Dr. Ed Roberts for necropsy procedures; Vinh Nguyen for overseeing pancreas dissociation; Matthias Hebrok, Andrei Goga, and Oakes and Papa Lab members for discussions.

