## Supplemental Figure 1 for "Parallel signaling through IRE1α and PERK regulates pancreatic neuroendocrine tumor growth and survival"

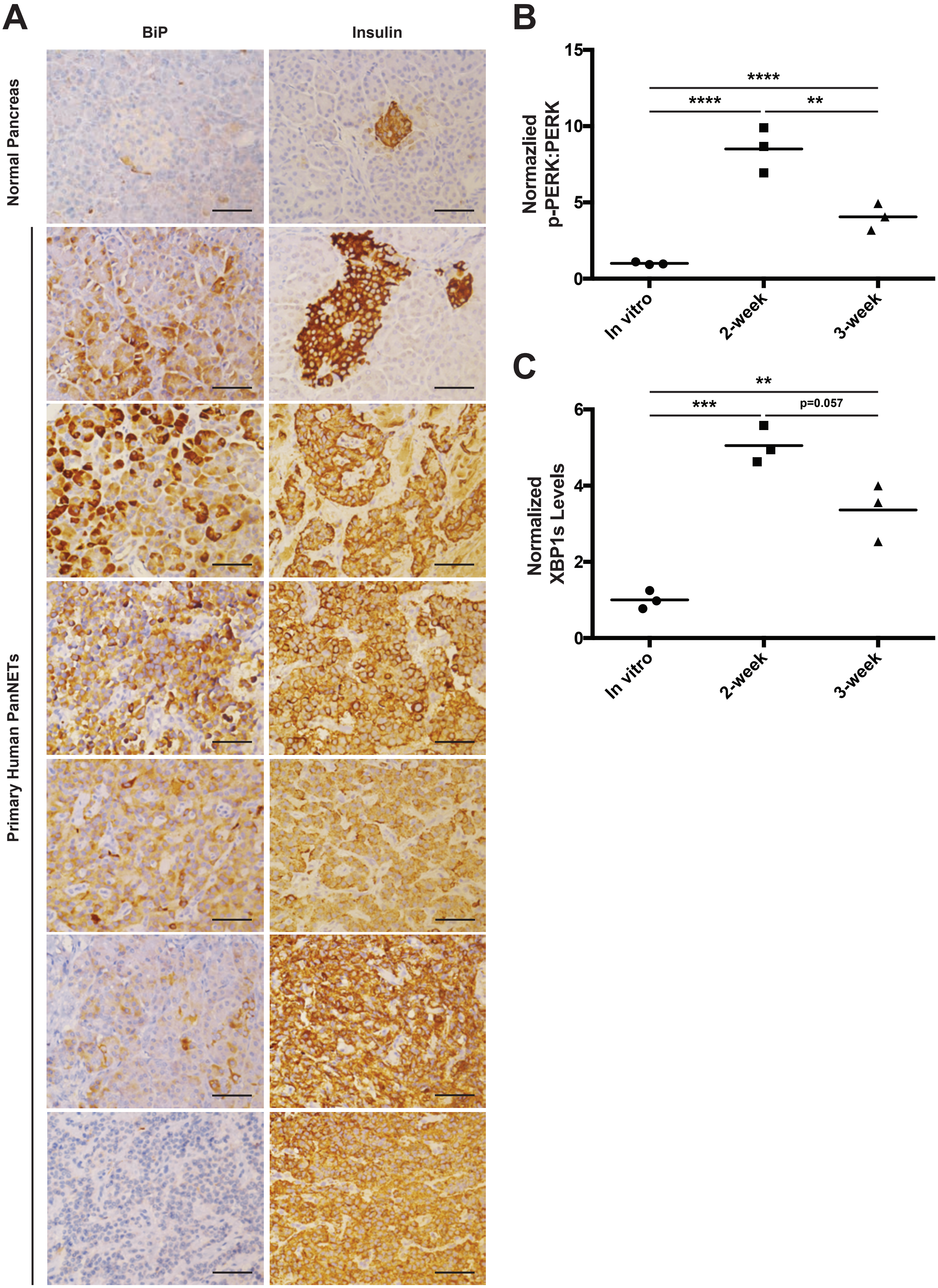


**Supplementary Figure S1. Majority of primary human PanNETs show elevated ER stress, Related to Figure 1**. A, IHC for BiP/GRP78 (left) and insulin (right) on normal human pancreas (top row) and a panel of six primary human PanNETs (Scale bars, 50 μm). B-C, Quantification of (B) the ratio of p-PERK to total PERK and (C) spliced XBP1 expression in Fig. 1L (n=3). **P<0.01, ***P<0.001, ****P<0.0001 (1-way ANOVA, Tukey test).
