## Supplemental Figure 2 for "Parallel signaling through IRE1α and PERK regulates pancreatic neuroendocrine tumor growth and survival"

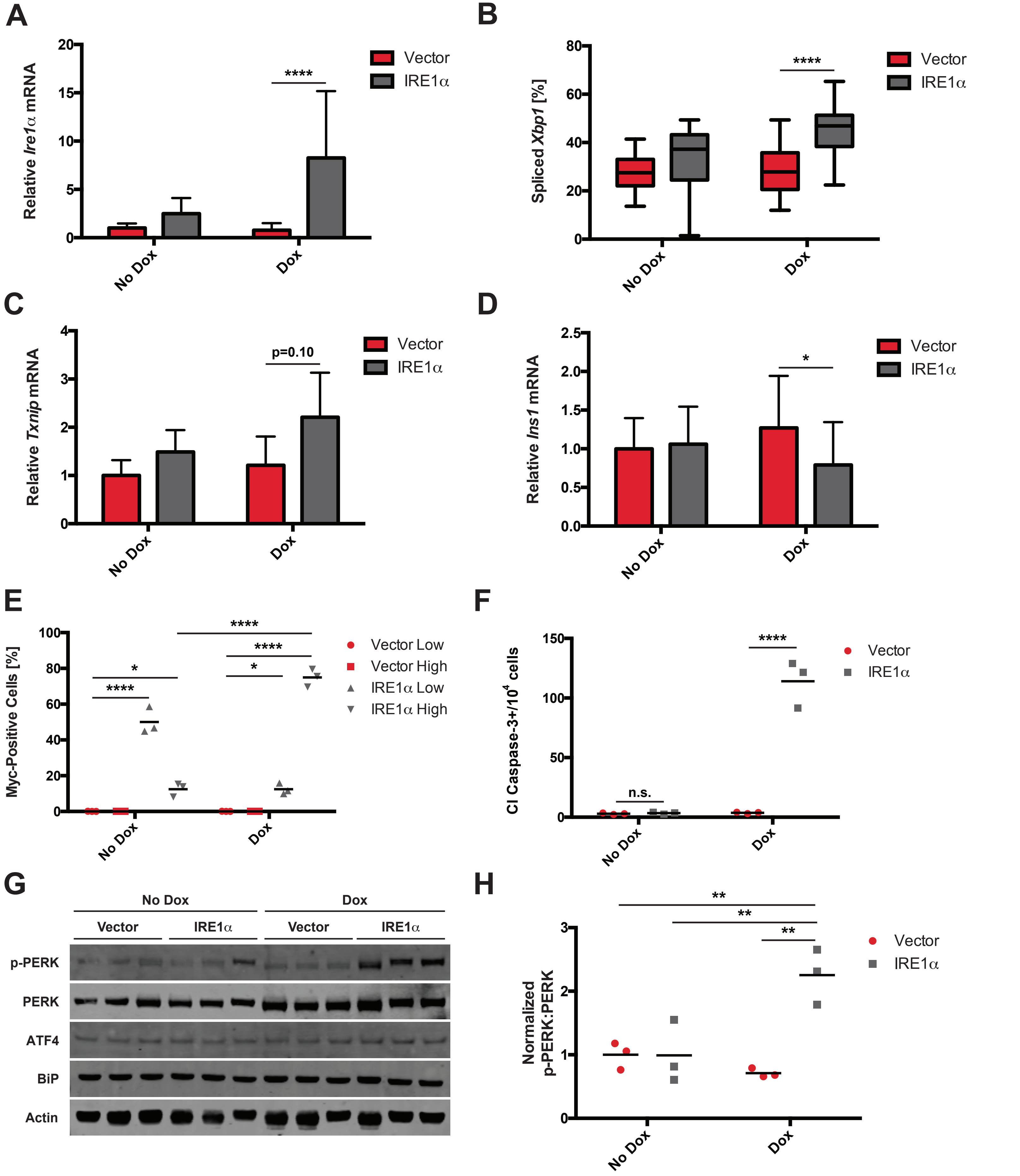


**Supplementary Figure S2. Manipulation of IRE1α adaptive vs. apoptotic signaling determines growth of INS-1 xenograft tumors, Related to Figure 2**. A, NSG mice were injected s.c. with INS-1 (vector) cells or INS-1 cells carrying a Dox-inducible, myc-tagged Ire1α gene (INS-1::Ire1α). Mice were fed regular or Dox chow as shown in Figure 1F, and Ire1α mRNA levels were quantified after 4 weeks (n ≥ 13). B, Percent spliced Xbp1 in INS-1 xenograft tumors at 4 weeks post-injection (n ≥ 13). C, mRNA levels of Txnip in INS-1 xenograft tumors treated +/- Dox for 96 hours at 2 weeks post-injection (n ≥ 5). D, mRNA levels of Ins1 in INS-1 xenograft tumors at 4 weeks post-injection (n ≥ 11). E-F, Quantification of (E) myc and (F) cleaved caspase-3 staining in Figure 2H and I (n=3). G, Immunoblots with the indicated antibodies of lysates from tumors in Figure 2H and I. H, Quantification of the ratio of p-PERK to total PERK in G (n=3). $*P<0.05$, $**P<0.01$, $****P<0.0001$ and n.s. = not significant (2-way ANOVA, Tukey test).
