## Supplemental Figure 3 for "Parallel signaling through IRE1α and PERK regulates pancreatic neuroendocrine tumor growth and survival"

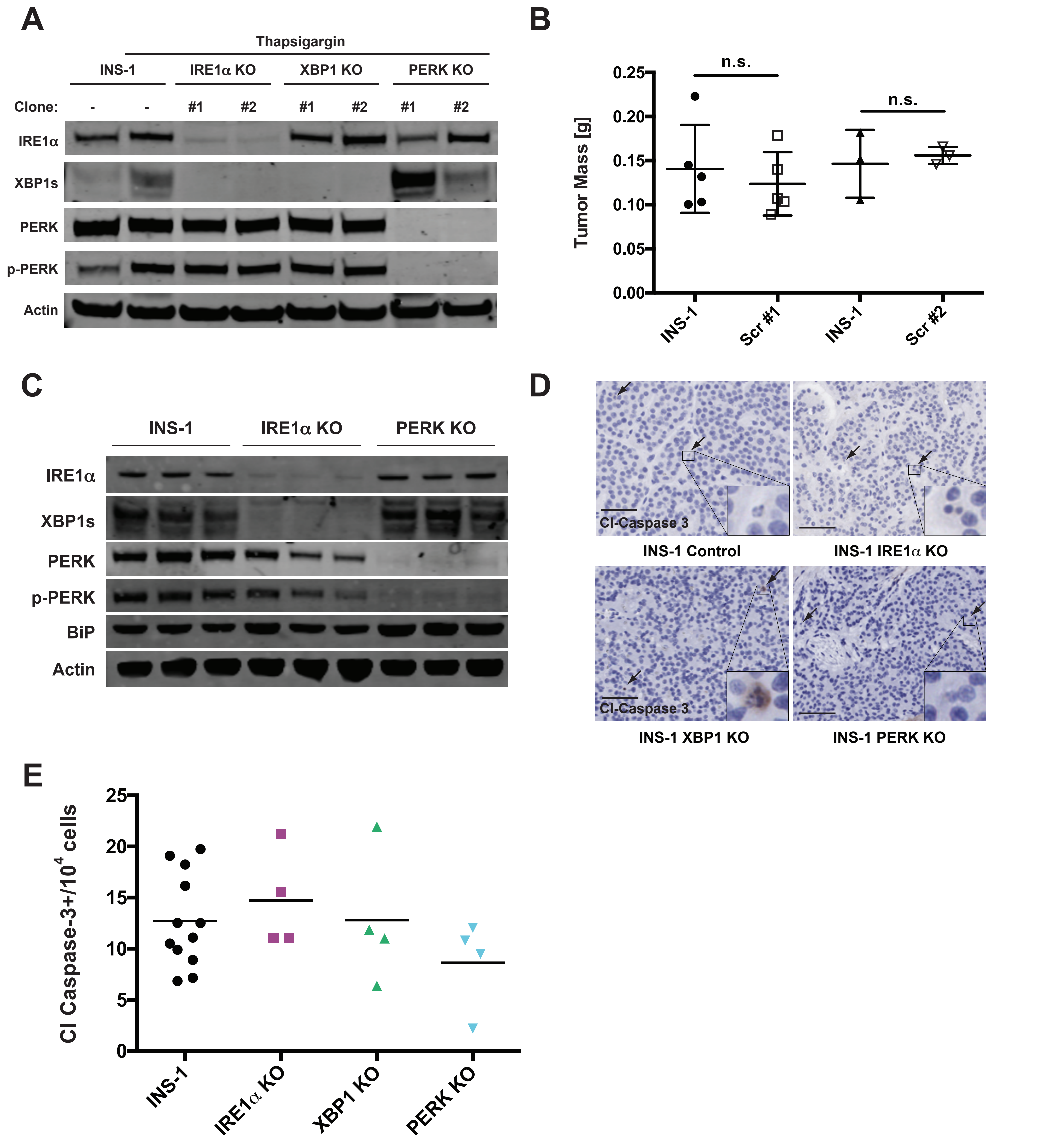


**Supplementary Figure S3. Scrambled CRISPR controls do not decrease INS-1 tumor burden, Related to Figure 3.** A, Cultured INS-1 control cells and the indicated CRISPR/Cas9 KO clones were treated +/- 31.25 nM thapsigargin for 3 hours prior to harvest to induce ER stress and then immunoblotted with the indicated antibodies. B, NSG mice were injected with INS-1 control and one of two unique scrambled (Scr) CRISPR control clones. Resulting tumors were harvested and weighed at 4 weeks post-injection (n ≥ 3). C, Immunoblots with the indicated antibodies of lysates from tumors in Figure 3C-E. D, Representative IHC for cleaved caspase-3 from control and indicated KO tumors at 4 weeks post-injection (scale bars, 50 µm). E, Quantification of cleaved caspase-3 staining in D (n ≥ 4). n.s. = not significant (1-way ANOVA, Tukey test).
