## Supplemental Figure 4 for "Parallel signaling through IRE1α and PERK regulates pancreatic neuroendocrine tumor growth and survival"

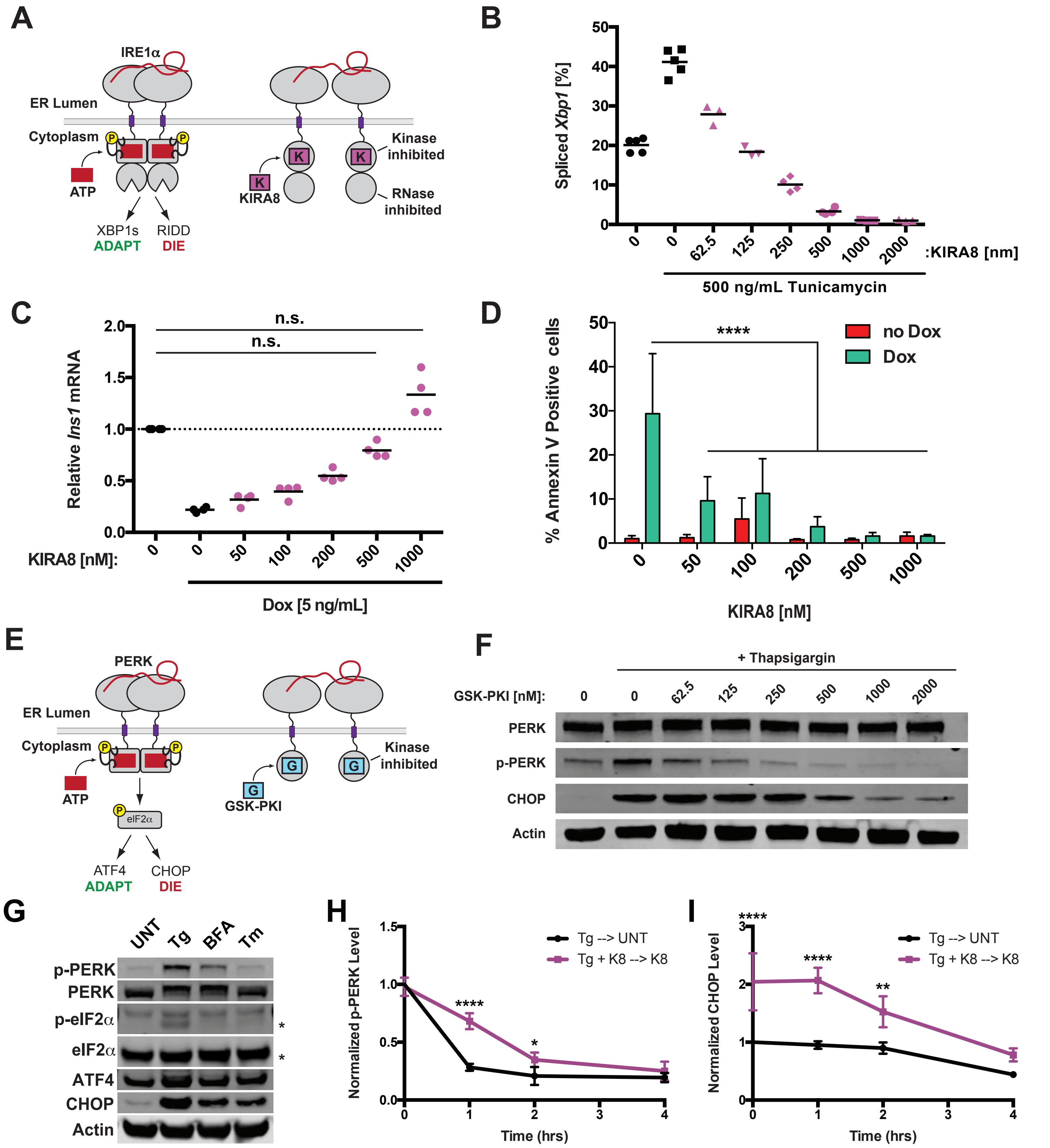


**Supplementary Figure S4. Pharmaceutical inhibitors of IRE1α and PERK are effective at inhibiting their kinase targets in vitro, Related to Figure 4**. A, Kinase Inhibiting RNase Attenuator 8 (KIRA8) binds directly to the kinase domain of IRE1α and allosterically inhibits the function of its RNase domain, thereby blocking IRE1α signaling in response to ER stress. B, Percent Xbp1 splicing from cultured INS-1 cells treated concurrently for 20 h with the indicated concentrations of tunicamycin and KIRA8 (n ≥ 3). C, mRNA levels of Ins1 in INS-1 IRE1α cells treated with the indicated concentrations of KIRA8 and Dox for three days (n = 4). D, Percentage of INS-1 IRE1α cells stained with Annexin V-FITC after three days of treatment with 10 ng/mL Dox and the indicated concentrations of KIRA8 (n = 8). E, GSK-PERK Kinase Inhibitor (GSK-PKI) binds directly to the kinase domain of PERK and blocks signaling in response to ER stress. F, Cultured INS-1 cells were treated concurrently for 16 h with 31.25 nM thapsigargin and the indicated concentrations of GSK-PKI, harvested and immunoblotted with the indicated antibodies. G, Immunoblots with the indicated antibodies of lysates prepared from INS-1 cells treated for 4 h with DMSO (UNT), 31.25 nM thapgsigargin (Tg), 500 ng/mL BFA, or 500 ng/mL tunicamycin (Tm). *Bottom band is eIF2α. H-I, Quantification of p-PERK (H) and CHOP (I) levels in Fig. 4F (n ≥ 3). *P<0.05, **P<0.01, ****P<0.0001, n.s. = not significant (1-way ANOVA, Tukey test in B-C; 2-way ANOVA, Tukey test in D; Sidak test in H-I).
