## Supplemental Figure 5 for "Parallel signaling through IRE1α and PERK regulates pancreatic neuroendocrine tumor growth and survival"

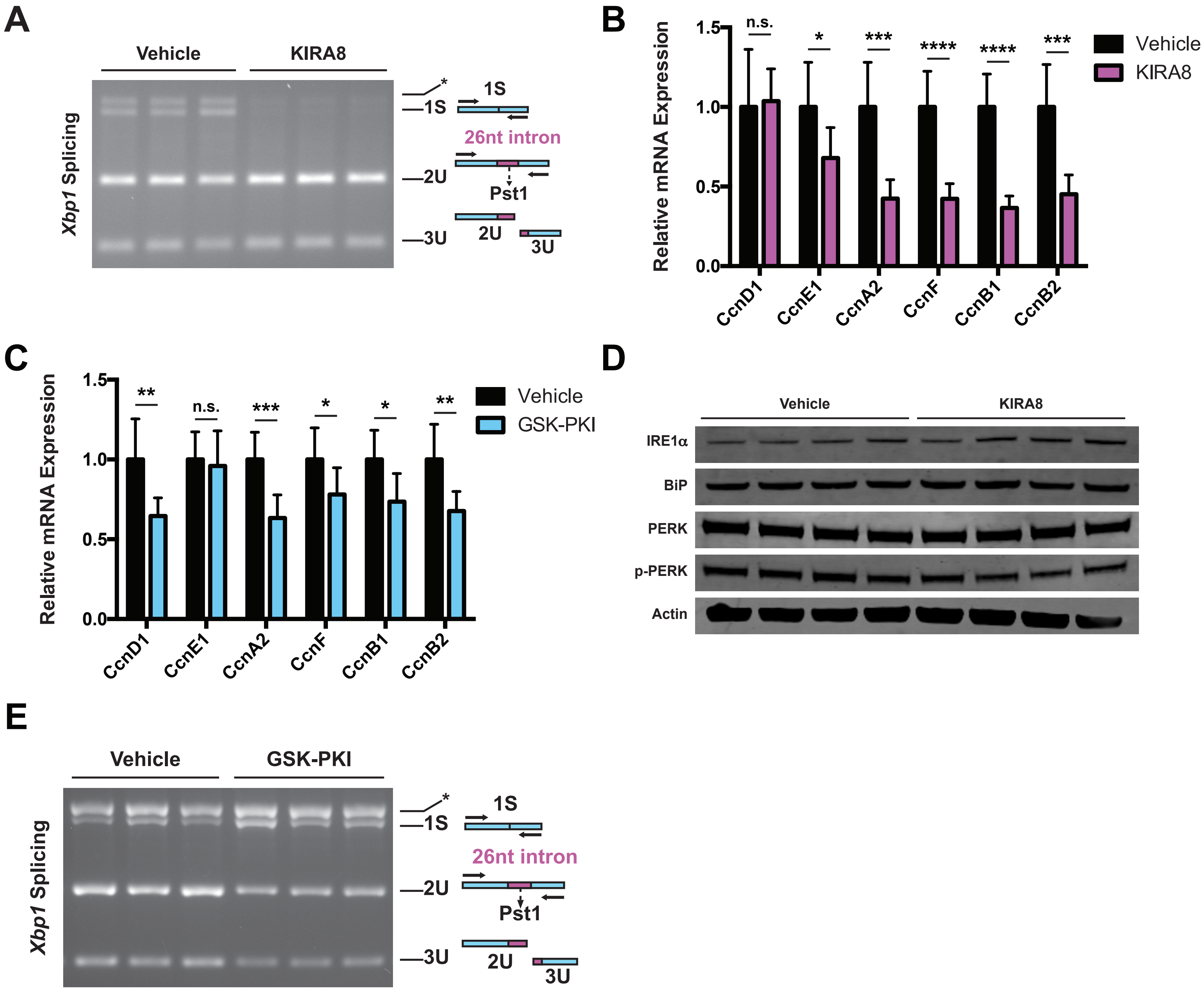


**Supplementary Figure S5. KIRA8 and GSK-PKI induce distinct apoptotic markers in INS-1 tumors, Related to Figures 5 and 6.** A, Representative DNA gel of the spliced (1S) and unspliced (2U and 3U) Xbp1 amplicons after PstI treatment. Quantified in Fig. 5A. B, Levels of the indicated mRNAs from tumors in Fig. 6A (n = 8). C, Levels of the indicated mRNAs from tumors in Fig. 6B (n = 8). D, Immunoblots with the indicated antibodies of tumors in Fig. 6A. E, Representative DNA gel of the spliced (1S) and unspliced (2U and 3U) Xbp1 amplicons after PstI treatment. Quantified in Fig. 6J. *P<0.05, **P<0.01, ***P<0.001, ****P<0.0001, n.s. = not significant (unpaired, two-tailed t tests).
