## Supplemental Figure 6 for "Parallel signaling through IRE1α and PERK regulates pancreatic neuroendocrine tumor growth and survival"

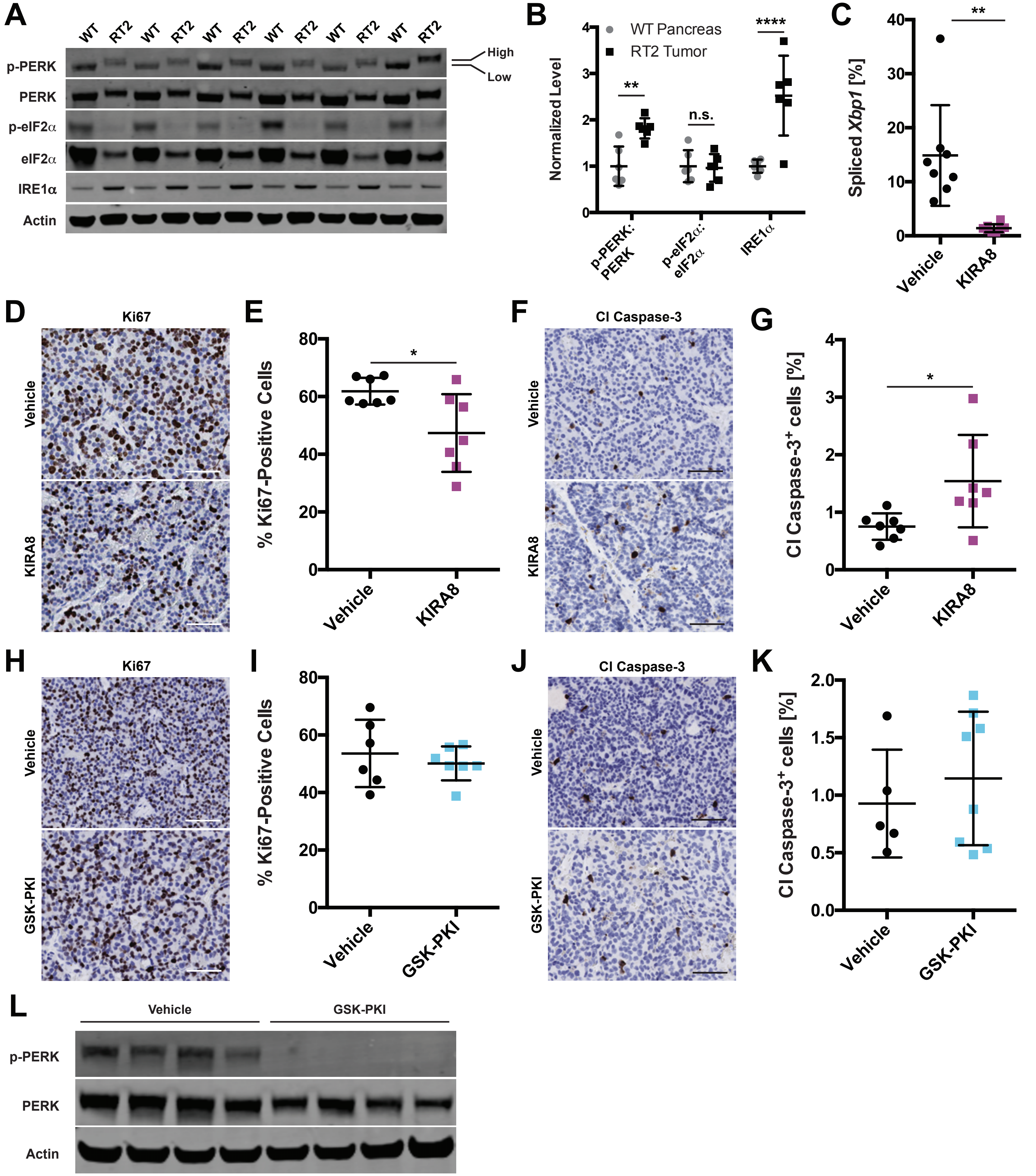


**Supplementary Figure S6. Effects of KIRA8 and GSK-PKI on RT2 tumors, Related to Figure 7.** A, Immunoblots with the indicated antibodies of lysates prepared from WT pancreata or RT2 tumors from 14-week-old mice. B, Quantification of the indicated targets in A (n=6). C, Percent spliced Xbp1 in RT2 tumors from 14-week-old mice treated 48 hours with vehicle or 50 mg/kg KIRA8. (n = 8). D, Representative IHC for Ki67 in tumors from C (scale bars, 50 μm). E, Quantification of Ki67 positive cells from D (n = 7). F, Representative IHC for cleaved caspase-3 in tumors from C (scale bars, 50 μm). G, Quantification of cleaved caspase-3 positive cells from F (n = 7). H, Representative IHC for Ki67 in tumors from 14-week-old RT2 mice treated 48 hours with vehicle or 50 mg/kg KIRA8 (scale bars, 50 μm). I, Quantification of Ki67 positive cells from A (n ≥ 6). J, Representative IHC for cleaved caspase-3 in tumors from A (scale bars, 50 μm). K, Quantification of cleaved caspase-3 positive cells from C (n ≥ 5). L, Immunoblots with the indicated antibodies of lysates from mice treated as in A. *P<0.05, **P<0.01, ****P<0.0001, n.s. = not significant (2-way ANVOA, Sidak test in B; unpaired, two-tailed t tests in C, E, G,I and K).
