## Supplemental Figure 7 for "Parallel signaling through IRE1α and PERK regulates pancreatic neuroendocrine tumor growth and survival"

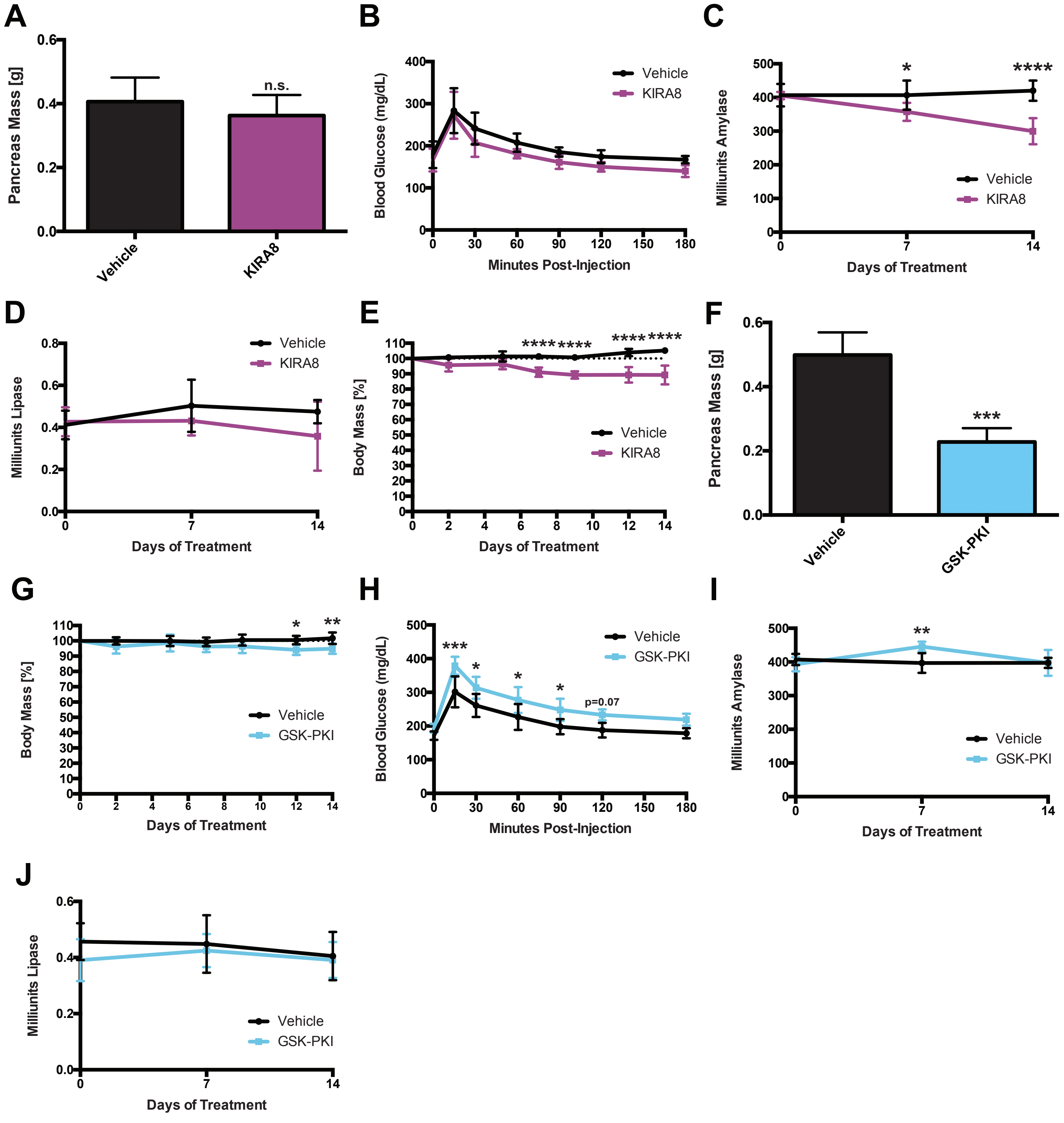


**Supplementary Figure S7. Effects of UPR blockade on C57BL/6 mice, Related to Figure 7.** A, Masses of pancreata from 14-week-old WT C57BL/6 mice after 14 days of treatment with vehicle or 50 mg/kg/d KIRA8 (n = 5). B, Mice treated as in A injected i.p. with glucose at t = 0. Blood glucose levels measured at the indicated time points (n = 6). C-D, Serum (C) amylase and (D) lipase levels of mice treated as in A (n = 6). E, Percent body mass at Day 0 of mice treated as in A (n = 6). F, Masses of pancreata from 14-week-old WT C57BL/6 mice after 14 days of treatment with vehicle or 50 mg/kg/d GSK-PKI (n ≥ 5). G, Percent body mass at Day 0 of mice treated as in F (n ≥ 5). H, Mice treated as in F injected with glucose at t = 0 and blood glucose levels measured at the indicated time points (n ≥ 5). I-J, Serum (I) amylase and (J) lipase levels of mice treated as in F (n ≥ 5). *P<0.05, **P<0.01, ***P<0.001, ****P<0.0001, n.s. = not significant (unpaired, two-tailed t tests in A and F; 2-way ANOVA, Sidak tests in B-E and G-J).
