## Supplemental Tables 1-3 for "Parallel signaling through IRE1α and PERK regulates pancreatic neuroendocrine tumor growth and survival"

Table S1. CRISPR/Cas9 guide Oligos

| Gene | Forward Sequence 5'-3' | Reverse Sequence 5'-3' |
|---|---|---|
| Ire1α #1 | caccGCAAGAGGACAGGCTCCATCAAG | aaacCTTGATGGAGCCTGTCCTCTTGC |
| Ire1α #2 | caccTGCCTGAACCAATTCCGGGA | aaacTCCCGGAATTGGTTCAGGCA |
| Perk #1 | caccAGATGGACGAATTGCCGCAC | aaacGTGCGGCAATTCGTCCATCT |
| Perk #2 | caccCGCGCGTGACTCCTGTTCGC | aaacGCGAACAGGAGTCACGCGCG |
| Scr #1 | caccGCACTACCAGAGCTAACTCA | aaacTGAGTTAGCTCTGGTAGTGC |
| Scr #2 | caccCCCCTTCGACCAGTCGGGTT | aaacAACCCGACTGGTCGAAGGGG |
| Xbp1 #1 | caccTTCCGGGCCCGCGAGCCGCA | aaacTGCGGCTCGCGGGCCCGGAA |
| Xbp1 #2 | caccGTTCCGGGCCCGCGAGCCGC | aaacGCGGCTCGCGGGCCCGGAAC |

Table S2. Genotyping/Sequencing Primers

| Gene | Forward Sequence 5'-3' | Reverse Sequence 5'-3' |
|---|---|---|
| Ire1α | TACAGGGCCATTTGAGGGAG | CGATCTCTCCAGCCCGAGTA |
| Xbp1 | GCCTGCAGGACCAATAAACG | GCGAATCTAACCCACCGTGA |
| Perk | GGAACCCTCGCTCAATGGG | CGAAACAATGAAAGCGGGGAA |

Table S3. qPCR Primers

| Species | Gene | Forward Sequence 5'-3' | Reverse Sequence 5'-3' |
|---|---|---|---|
| Human | ATF4 | GTTCTCCAGCGACAAGGCTA | ATCCTCCTTGCTGTTGTTGG |
| Rat | Actin | GCAAATGCTTCTAGGCGGAC | AAGAAAGGGTGTAAAACGCAGC |
| | Bim | CGGATCGGAGACGAGTTCAA | TAACCATTTGCGGGTGGTCT |
| | CcnA2 | AGTGCCGCTGTCTCTTTACC | GGGGTGATTCAAAACTACCATCC |
| | CcnB1 | TGAACTTCAGTCTGGGTCGC | TGAACTTCAGTCTGGGTCGC |
| | CcnB2 | TGGCTGGTCCAAGTCCATTC | GAGCTGTGATCCCAACCAGT |
| | CcnD1 | TCAAGTGTGACCCGGACTG | GGATCGATGTTCTGCTGGGC |
| | CcnE1 | CGGACACAGCTTCGGGTCT | TCTGGATCGGACTGAGAGGT |
| | CcnF | CCTCCCTCTGGCACTTTGAG | CGGGCAAAGAATGGCCTAGA |
| | Gapdh | CAGGGCTGCCTTCTCTTGTG | AACTTGCCGTGGGTAGAGTC |
| | Ins1 | GTCCTCTGGGAGCCCAAG | ACAGAGCCTCCACCAGG |
| | Ins2 | GGGAGCGTGGATTCTTCTACAC | CCACTTGTGGGTCCTCCACTT |
| | Ire1α | GTCCCACTTTGTGTCCAATGG | TCCCCAGACATGAAGGTCA |
| | Txnip | CTGATGGAGGCACAGTGAGA | CTCGGGTGGAGTGCTTAGAG |
